# Where you aim – not how you aim – affects implicit recalibration in visuomotor adaptation

**DOI:** 10.64898/2026.02.19.706918

**Authors:** Yiyu Wang, Jordan A. Taylor

## Abstract

The influence of explicit strategies on implicit recalibration during visuomotor adaptation has become a central question in motor learning. Because the two systems operate in tandem, explicit strategies could indirectly influence implicit recalibration. However, explicit strategies are not unitary: they may rely on algorithmic-based computations or memory-based retrieval of cached solutions. This raises the possibility that different strategy implementations interact with the cerebellar-based implicit recalibration system in qualitatively distinct ways, especially given that these strategies likely rely on different frontoparietal networks. Here, we tested whether the type of explicit strategy modulates implicit recalibration. Across a set of experiments, we observed subtle differences in the spatial profile of implicit generalization: the algorithmic strategy produced a broader generalization pattern than the retrieval strategy, even after controlling for intertrial decay, generalization structure, and between-target interactions. While this pattern is suggestive of greater flexibility afforded by algorithmic strategy use compared to memory-based retrieval, it could instead arise from increased variability in explicit aiming, which constitutes the input data driving implicit recalibration. Indeed, when we isolated the direct contribution of each strategy to implicit recalibration by rigorously controlling for reach variability and using error-clamp feedback to ensure uniform implicit learning conditions, we found no difference in implicit recalibration across strategies. Together, these findings suggest that while algorithmic and retrieval strategies differ in their behavioral signatures and influence the movement plan, the implicit recalibration process itself remains rigid with respect to the strategy employed.

## Introduction

The interaction between explicit and implicit processes has recently become a central focus in visuomotor adaptation, as conflicting findings across studies have left the precise influence of explicit processes on implicit recalibration unresolved. On the one hand, some studies support the view that explicit and implicit processes operate largely independently (Bond and Taylor, 2015; McDougle et al., 2015; Morehead et al., 2017), as reflected in their distinct learning signatures and their reliance on different neural systems (Mazzoni and Krakauer, 2006; Taylor and Ivry, 2011, 2014; Butcher et al., 2017; Areshenkoff et al., 2024). On the other hand, several studies suggest that explicit strategies may directly (Albert et al., 2022) or indirectly influence implicit recalibration (Day et al., 2016; McDougle et al., 2017). Evidence for such interactions includes plan-based generalization, in which implicit recalibration is centered on the explicit aiming direction (Day et al., 2016; McDougle et al., 2017), competition for a shared error signal (Albert et al., 2022), and nonlinear patterns of interaction revealed through various methodological approaches (Maresch et al., 2021; t Hart et al., 2024; Chen and Taylor, 2025).

These mixed findings may arise from subtle differences in task design, which can shift the nature of the explicit strategies participants employ (McDougle and Taylor, 2019; Velazquez-Vargas and Taylor, 2024); such shifts, in turn, can modulate the expression of implicit recalibration in distinct ways (Albert et al., 2022). At least two distinct types of explicit strategies have been identified in visuomotor adaptation. The first is an algorithmic strategy, in which participants mentally rotate their intended movement vector away from the target by an amount equal in magnitude and opposite in direction to the imposed cursor rotation (Georgopoulos et al., 1986; Georgopoulos and Pellizzer, 1995; Sanes and Donoghue, 2000; McDougle and Taylor, 2019). This visuomotor mental-rotation strategy is highly flexible, but it comes with a cost: preparation time increases linearly with the size of the required rotation, paralleling the classic mental-rotation effects (Shepard and Metzler, 1971). In contrast, once a successful solution has been identified, participants may instead rely on a retrieval strategy, in which the previously successful aiming solution is cached in working memory and retrieved at minimal computational cost (McDougle and Taylor, 2019; Fresco et al., 2023; Velazquez-Vargas and Taylor, 2024). However, this retrieval process is constrained by its reliance on previously learned stimulus–response associations (Logan, 1988). As a result, retrieval-based strategies show limited generalization and tend not to extend far beyond the specific conditions experienced during training (McDougle and Taylor, 2019).

A common method for manipulating strategy use in visuomotor rotation tasks is to vary the number of training targets (set size; (McDougle and Taylor, 2019)). A high set size overwhelms working memory, rendering retrieval strategies ineffective (McDougle and Taylor, 2019; Velazquez-Vargas and Taylor, 2024; Bejjanki and Taylor, 2026) and forcing reliance on an algorithmic strategy. Previous research has demonstrated that manipulation of set size appears to impact the magnitude of cerebellar-dependent implicit recalibration (Bond and Taylor, 2015; Neville and Cressman, 2018) and mediates the performance of patients with cerebellar-ataxia (Gibo et al., 2013; Tsay et al., 2022; Hadjiosif et al., 2023).

The effect of set size on implicit recalibration could be the result of temporal decay, which is supported by prior work finding that implicit recalibration comprises temporally persistent and temporally volatile components(Hadjiosif et al., 2023). As set size increases, the interval between successive visits to the same target grows, allowing the volatile component to decay before it can carry over and accumulate on the next encounter. This rapid loss would reduce net trial-to-trial buildup, yielding smaller apparent implicit recalibration at larger set sizes (Hadjiosif et al., 2023; Hadjiosif et al., 2024). Alternatively, it could be that different strategies, which are guided by set size (Miller, 1956; Oberauer et al., 2016; Cowan, 2017) and draw upon different frontoparietal networks (D’Esposito, 2007; Engelhardt et al., 2019; Reineberg et al., 2022), lead to different interactions with the cerebellum, which, in turn, reshape the profile of implicit recalibration (Ravizza et al., 2006; Buckner et al., 2011; Bernard and Seidler, 2013).

In this study, we sought to systematically test whether algorithmic and retrieval strategies differentially shape implicit recalibration during visuomotor adaptation. To obtain an unbiased estimate of implicit learning and isolate the unique contribution of each strategy, we conducted three experiments that progressively tightened control over key confounding factors. These included: plan-based generalization, which can broaden the implicit generalization function due to variability in movement planning, and inter-trial interval, which may produce unequal levels of adaptation between groups through differential memory decay. We hypothesized that the strategy engaged during adaptation could affect both the magnitude and generalization profile of implicit recalibration. In particular, we predicted that relying on an algorithmic strategy—requiring computation on each trial—would lead to a broader implicit generalization function compared to repeatedly retrieving a well-learned solution.

## Results

### Experiment 1: Does the use of either an algorithmic or a retrieval strategy result in different implicit generalization?

Experiment 1 was designed to test whether implicit recalibration differs depending on whether adaptation is supported by an algorithmic or retrieval-based strategy. Participants completed a visuomotor rotation task and were assigned to one of two conditions that biased strategy use (Figure 1A). In the algorithmic condition, rotated feedback was provided across all training targets to encourage trial-by-trial computation of the aiming solution (Figure 1B). In the retrieval condition, rotated feedback was restricted to a single Critical target, with brief pretraining at that location to encourage retrieval of a cached solution (Figure. 1C). Importantly, the arrangement of the Critical target and Non-Critical targets were specifically designed to limit rotation training at locations neighboring the Critical target (Figure 1B – C). Implicit recalibration was assessed with Exclusion trials (Werner et al., 2015; Maresch et al., 2021), in which participants were instructed to withhold the strategy and reach without feedback to probe targets surrounding the Critical target (Figure 1D). These probe trials allowed us to estimate both the magnitude of implicit recalibration and its generalization around the Critical target location periodically throughout training (Figure 1E)

**Figure 1.**
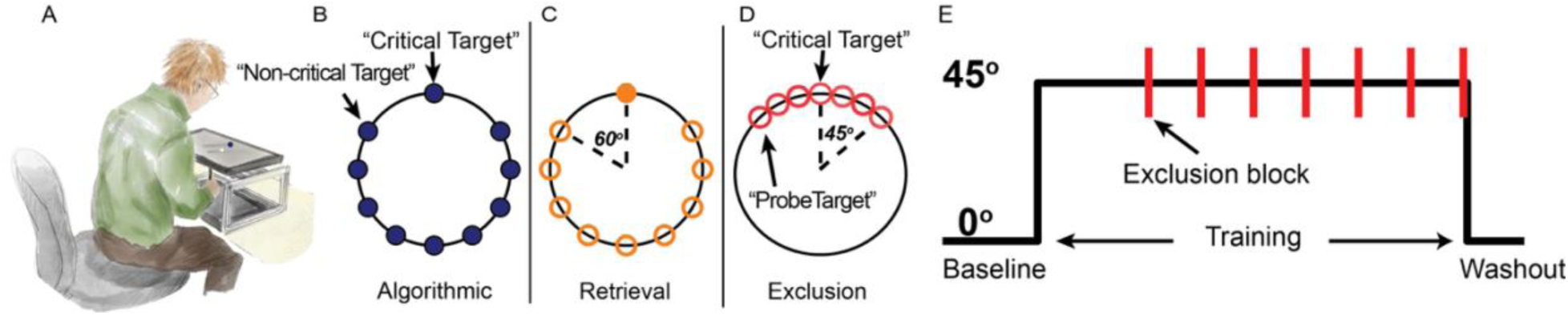
Experimental setup and procedures. A) Setup for the experiment. B) Illustration of experimental workspace for the Algorithmic group (blue) and C) the Retrieval group (orange) during training. A 45° visuomotor rotation was applied to the Critical target and all Non-Critical targets (filled) for the Algorithmic group, while no rotation was applied to the Non-Critical targets for the Retrieval group (empty). The 60° gap was applied to separate the space between the critical and Non-Critical targets. D) Probe targets (empty) neighbored the Critical target. No feedback was provided during Exclusion trials and participants were instructed to withhold strategy use. E) No rotation was applied during baseline (80 trials), a rotation was applied during training (356 trials), Exclusion trials (7 trial mini-blocks) were intermittently dispersed during training (49 Exclusion trials in total), and cursor feedback was withheld during the washout block at the end of training (48 trials).

#### Adaptation performance

Both groups had little difficulty following instructions to reach the targets directly during the baseline phase, as reflected by no difference in error (t(84) = -0.192, p = 0.848). When the 45° visuomotor rotation was introduced during the training phase, both groups adapted quickly at the Critical target location, with reach angles increasing from early to late training and approaching near-complete compensation by the end of training (Figure 2A, see Figure 1A in Supplemental Information (Figure S1A) for complete trial-by-trial time courses). At Non-Critical targets, errors remained elevated over the course of training for the Algorithmic group since the task design demanded they compensate for the rotation, while the Retrieval group did not (Figure 2B, see Figure S1B for trial-by-trial hand angle time courses). To determine whether there were any differences in the learning function between the groups, given the differences in task demands, the hand angles were analyzed using a linear mixed-effects model with Group (algorithmic and retrieval) and Time (early and late) as factors. As expected, we found a main effect of Time (b = - 1.25, SE = 0.33, t(168) = -3.76, p < .001), suggesting a continuous improvement in reach performance over the course of training. We also found a main effect of Group (b = -0.96, SE = 0.43, t(168) = -2.23, p = .003), suggesting a distinct learning pattern associated with each strategy. More interestingly, we found an interaction between Group and Time (b = –0.74, SE = 0.33, t(168) = –2.23, p = .027), with linear contrast analyses revealing that the Retrieval group showed faster learning than the algorithmic group. Specifically, during the early phase of training, the retrieval group (42.21 ± 4.62°) exhibited significantly greater hand angles than the algorithmic group (38.82 ± 8.44°; p = .002), but by the end of training, both groups achieved similar levels of performance (Retrieval 43.23 ± 2.12°; Algorithmic 42.81 ± 2.59°, p = .695). Within-group comparisons indicated that the degree of adaptation in the Retrieval group remained stable across training, with no significant difference between the early and late phases (p = .28). This finding suggests that brief exposure to adaptation at the Critical target location before training was sufficient to cache the correct aiming strategy in the retrieval group, thereby minimizing reliance on an algorithmic strategy during early adaptation and reducing a potential confound in interpreting implicit recalibration at the Critical target location. In contrast, the degree of adaptation reflected by hand angle increased from the early to late phase of training in the algorithmic group (p = 0.002). At a minimum, these findings suggest that storing and retrieving a memory from a short-term memory cache confers more rapid performance improvements than executing an algorithmic strategy – a point that is supported by decreased reaction time for the retrieval group (see *Preparation Time* below).

**Figure 2.**
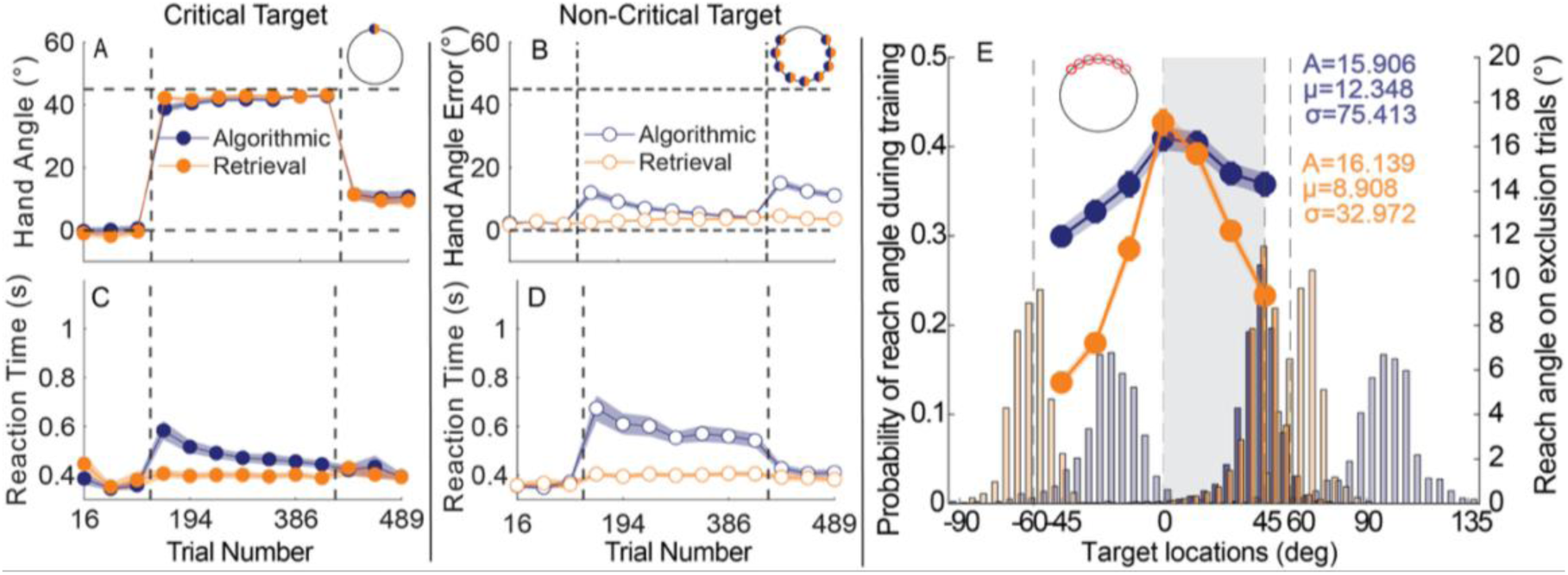
Adaptation performance, reaction time, and implicit generalization functions for the algorithmic and retrieval strategies in Experiment 1, with potential confounds controlled relatively loosely. A) Adaptation performance measured at the Critical target location, reflected by the hand angles reaching near 45°. Vertical dashed lines indicate onset and offset of the training phase. Each data point represents binned hand angles for one cycle of training (see supplementary material for complete time course data). A mini workspace diagram is included in each figure to show the target locations where data are demonstrated. B) Absolute hand angle error (unsigned error) is calculated for all Non-Critical target locations to reflect the accuracy of performance. Given that each group had distinct task instructions at the Non-Critical target, absolute hand angle error can ensure a numerical comparison. C) Reaction time for the Critical target and D) Reaction time for Non-Critical target locations. E) Two generalization curves (right y-axis) for the algorithmic (blue) and retrieval (orange) groups are shown across the corresponding probe target locations (x-axis, -45° – 45°). Beneath these curves, bar plots illustrate the distribution of reach directions for each group (left y-axis), with darker colors indicating reaches to the Critical target and lighter colors representing reaches to adjacent Non-Critical target locations (±60°). Note that implicit recalibration can be acquired within the area containing the blue or light blue bars due to the use of continuous feedback; in contrast, no feedback was shown for the reaches represented by the lighter orange bar in the retrieval group, which would not produce the implicit recalibration. The shaded gray region marks the “aiming zone,” where participants were most likely to aim when counteracting the 45° rotation at the Critical target. The amplitude, mean, and width parameters displayed in the figure were derived from Gaussian function fits.

At Non-Critical targets, we sought to confirm that participants consistently followed the task instructions, as performance at these targets was critical for reinforcing the intended strategy (algorithmic vs. retrieval). Because the groups received different instructions at these locations—the algorithmic group continued adapting to the 45° rotation, whereas the retrieval group aimed directly at the target—we quantified reaching accuracy to make performance numerically comparable across groups (Figure 2B, see Figure S1B for trial-by-trial hand angle time courses). The analysis revealed that accuracy for the algorithmic group progressively increased from early to late training to offset the visuomotor rotation (b = -7.88, SE = 0.98, t (168) = -8.01, p < .001), ultimately, diminishing reach error to 4.14 ± 2.32° from 12.02 ± 9.79° during the Early training (Figure 2B, Figure S1B). In contrast, because they were instructed to aim directly toward these Non-Critical targets and no rotation was applied, the hand angles of the retrieval group remained stable across training (Early: 2.54 ± 2.44°, Late: 3.92 ± 2.84°; p = .163). This trial-dependent accuracy pattern at the Non-Critical target locations closely mirrored that at the Critical target location, indicating that participants followed the instructions, which is required for successfully implementing the intended strategy across groups.

#### Preparation time

Preparation time, operationalized here as RT, provides a window for identifying the strategic processes underlying visuomotor adaptation (McDougle and Taylor, 2019). Despite both groups achieving similar performance at the Critical target, there were clear differences in their RT (Figure 2C, Figure S1C for the trial-by-trial RT time courses). Across training trials, the algorithmic (p < 0.01) and retrieval groups (p = 0.04) showed increased RT compared to their baseline levels, indicating the implementation of cognitive strategies (Figure S1E, F for comparing mean RT between baseline and training phases). Indeed, a linear mixed-effects model examining reaction time revealed a significant main effect of Group (b = 0.058, SE = 0.014, t(168) = 4.25, p < .001), indicating that the retrieval group responded faster than the algorithmic group across training. There was also a significant main effect of Time (b = 0.04, SE = 0.008, t(168) = 4.57, p < .001), showing that reaction times decreased from early to late practice; More interestingly, the group × time interaction was significant (b = 0.03, SE = 0.008, t (168) = 3.56, p < .001) with only the algorithmic group showing improvements in RT over the course of training (Early: 0.58 ± 0.25 s; Late 0.44 ± 0.10 s; p < .001). In contrast, the retrieval group maintained stable reaction times from early (0.41 ± 0.11 s) to late (0.39 ± 0.08 s) (p = .478). At the Critical target location, this pattern of results supports the interpretation that participants in the retrieval group cached a strategy in working memory, while the algorithmic group relied primarily on mental-rotation strategies (McDougle and Taylor, 2019; Velazquez-Vargas and Taylor, 2024).

Further validation that our experimental design effectively guided the two groups toward distinct strategies is provided by the RT data at the Non-Critical target locations (Figure 2D, Figure S1D). The successful implementation of these strategies depended on participants closely following the instructions at the Non-Critical targets. We found a significant main effect of Group (b = 0.10, SE = 0.02, t (168) = 5.83, p < .001), Time (b = −0.03, SE = 0.01, t(168) = −2.62, p = .01), and interaction (b = 0.03, SE = 0.01, t (168) = -2.67, p = .008). Planed linear contrast showed that RT in the retrieval group remained stable across training trials (p = .97), whereas RT in the algorithmic group became faster during late training than during early training (p < .01). The RT pattern at the Non-Critical target locations mirrored that observed at the Critical target location, confirming that participants in the algorithmic group relied on a visuomotor mental rotation strategy across all targets, both Critical and Non-Critical.

#### Implicit recalibration

The primary goal of this study was to determine if different kinds of explicit strategies have downstream consequences for implicit recalibration, in particular at the Critical target. To quantify implicit recalibration in relative isolation of strategies, we periodically leveraged Exclusion probe target trials around the Critical target. Here, participants were instructed to refrain from using any strategies they may have developed and instead simply reach directly to the presented target without cursor feedback. Because plan-based generalization can shift the center of the implicit generalization function, these Exclusion trials were drawn from -45° to 45° centered around the Critical target to increase the likelihood of capturing the true amplitude of the implicit recalibration (Day et al., 2016; McDougle et al. 2017, Poh and Taylor 2019).

Prior work has shown that implicit recalibration is reduced when adaptation must be learned for multiple targets, raising the possibility that strategy use may differentially shape implicit generalization (Bond and Taylor, 2015). To test this idea, we first compared hand angles during Exclusion trials at the Critical target and observed no difference in the magnitude of implicit recalibration between retrieval (17.09 ± 9.51°) and algorithmic strategy (16.37 ± 9.09°, t(84) = - 0.47, p = 0.64) at the Critical target probe location (Figure 2E). Inspection of the full generalization function indicated that strategy type did not affect the overall amplitude of implicit recalibration (Figure 2G and Figure S5A for model fitting, algorithmic: 15.91; retrieval: 16.14; bootstrap, p = 0.42). Likewise, the center of the generalization function did not differ between groups (Figure 2, algorithmic: 12.35; retrieval: 8.91; bootstrap, p = .35). At first glance, these results suggest that strategy type does not influence implicit recalibration. However, a different pattern emerged when we examined the breadth of the generalization function. The generalization function was significantly broader for the algorithmic group (σ = 75.41°) than for the retrieval group (σ = 32.97°) (Figure 2G, bootstrap, p < .001), indicating that strategy type selectively modulated the spatial extent of implicit generalization.

While this difference in the breath of the generalization function points to the potential influence of strategy use on the operation of implicit recalibration, several other first-order factors could potentially explain the differences observed on Exclusion trials: 1) unequal adaptation magnitude prior to the assessments of implicit recalibration, 2) incomplete strategy withholding during Exclusion trials, and 3) differences in reach angle variability between the algorithmic and retrieval groups. First, we administered a “top-up” trial at the Critical target location prior to each Exclusion assessment to ensure that the levels of adaptation were similar between groups, as well as minimize the effects of temporal decay (Hadjiosif et al., 2023). Indeed, analysis of hand angles across all “top-up” trials revealed no significant main effect of Group (F (1, 575) = 0.19, p = .66), indicating that overall adaptation did not differ between the two groups across top-up trials. There was also no significant main effect of the trial (F (6, 575) = 1.41, p = .24), showing that adaptation levels remained stable across repeated top-up trials. Critically, the group × trial interaction was not significant (F (6, 575) = 0.37, p = .90), indicating that the two groups exhibited highly similar patterns of adaptation across all seven blocks (Figure S4A for “top-up” trial performance across block).

Second, we ruled out the possibility that the use of distinct strategies during the Exclusion block contributed to the observed differences in the generalization function. More specifically, in the algorithmic group, although RT during the Exclusion block (0.40 ± 0.09 s) was significantly slower than baseline RT (0.35 ± 0.06 s, p < .01), it was significantly faster than RT during training (0.58 ± 0.20 s, p < .01). Together, these results suggest that participants attempted to suppress strategic processing during the Exclusion block, consistent with the instruction to aim directly at the probe target locations. The fact that RT remained elevated relative to baseline likely reflects a task-switch cost associated with switching from a training task that required greater computational demands to compensate for the 45° rotation to an Exclusion task that required only direct aiming to the target (Figure S1E). In the retrieval group, RT during the Exclusion block (0.38 ± 0.08 s) did not differ from baseline RT (0.37 ± 0.12 s, p = .99) but was significantly faster than RT during training (0.41 ± 0.07 s, p < .01, Figure S1F), supporting that participant attempted to minimize the use of strategy in the retrieval group. When comparing RTs across groups, the analysis revealed no overall difference between groups during the Exclusion block (F(1, 588) = 1.07, p = .30), suggesting that participants in the algorithmic group did not engage in the high-cost mental-rotation strategy typically associated with slower responses. This result also serves as a manipulation check, indicating that participants in both groups followed the instruction to withhold their strategies. Although reaction times exhibited small fluctuations across the seven blocks (main effect of trial: F (6, 588) = 4.92, p < 0.01), importantly, they did not differ between groups (trial × group interaction: F (6, 588) = 1.17, p = .32, Figure S1C-D, Figure S4D for Exclusion block RTs comparison across groups).

Finally, upon closer inspection, the distribution of reaches during training were not identical between the groups as the algorithmic group produced more variability than the retrieval group (Figure 2E; bootstrap, p < .001). According to the plan-based generalization account (McDougle et al., 2017), the greater variability in the distribution of reaches in the algorithmic group suggests more variable re-aiming plans in the training workspace, which may have broadened the generalization function. In particular, the spillover of implicit recalibration in this experiment may have arisen from two sources: (1) the generation of re-aiming plans for the non-critical targets, and (2) greater variability in the re-aiming plans for the critical target. Supporting this interpretation, we found that implicit recalibration at the Non-Critical target locations was acquired only in the algorithmic group, where online feedback was provided at those locations, as reflected by larger aftereffects at the Non-Critical targets in the algorithmic group than in the retrieval group. (algorithmic: 15.00 ± 8.36°; retrieval: 4.59 ± 3.05°), t(84) = 7.68, p < .001; Figure S1G, H). The minimal aftereffects in the retrieval group can be anticipated because no visual feedback was provided at the Non-Critical target locations (Figure 1C). Accordingly, the shape of the implicit generalization function around the Critical target in each group was consistent with the extent of implicit recalibration at the Non-Critical target locations. Moreover, the variability of the reach distribution at the Critical target location was significantly greater in the algorithmic group than the retrieval group (bootstrap, p < 0.01, Figure S6A). This may further contribute to the broader breadth of the generalization function observed in the algorithmic group. Together, these findings suggest that the broader generalization observed in the algorithmic group may have been driven, at least in part, by spillover learning from adjacent targets and the greater variability in reach. Thus, Experiment 1 did not provide sufficient evidence to determine whether algorithmic and retrieval strategies differentially affect implicit recalibration.

### Experiment 2: Is the increased breadth of implicit generalization the result of “error spill over”?

To address the key confounds identified in Experiment 1, we designed Experiment 2 to suppress error spillover more directly and better isolate implicit recalibration at the region where the probe targets are located. Specifically, we increased the spatial separation between the Non-Critical targets and Critical target, and introduced delayed endpoint feedback at all Non-Critical locations for both groups to ensure equal comparison. Continuous feedback was retained at the Critical target only to permit implicit recalibration, enabling us to measure the influence of strategies on implicit recalibration. Because delayed endpoint feedback is known to significantly blunt implicit recalibration (Brudner et al., 2016), these modifications were intended to minimize implicit recalibration spillover from the Non-Critical targets (Figure 3A,B). Apart from these changes, the experimental protocol was identical to that of Experiment 1. The primary goal of Experiment 2 was to build on Experiment 1 by testing whether algorithmic and retrieval strategies produce different levels of implicit recalibration under stricter experimental control.

**Figure 3.**
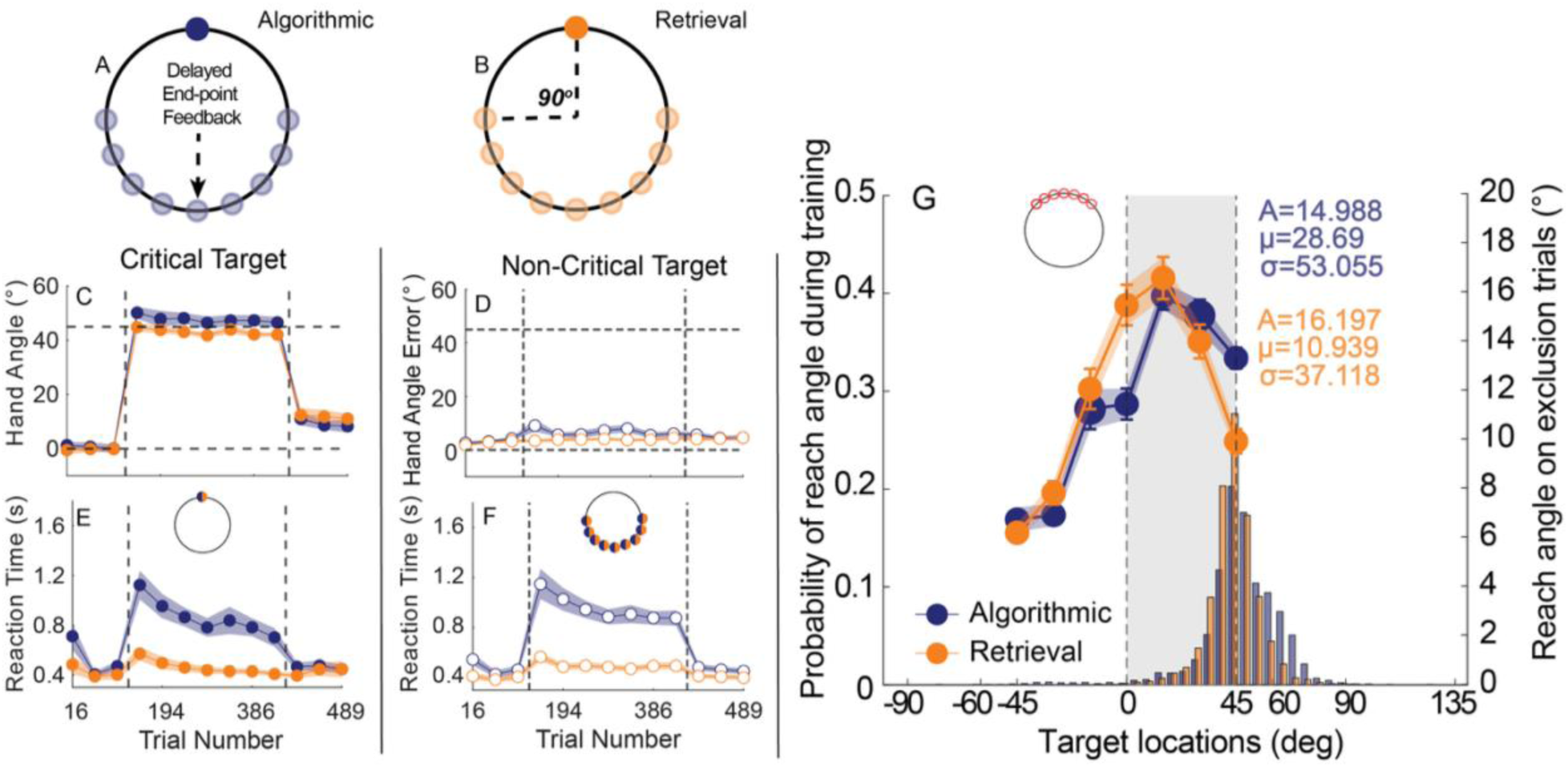
A stricter control narrows down the implicit generalization function in the algorithmic group. A) Diagram of workspace setup and target locations for algorithmic strategies in Experiment 2. Participants receive delayed endpoint feedback at all Non-Critical targets, so that implicit recalibration at all Non-Critical targets should not occur. The continuous online feedback was applied to the Critical target, enabling implicit recalibration. The Critical target is selected at least ±90° away from Non-Critical targets to further attenuate the influence of aiming for adapting 45° rotations at the Non-Critical targets. B) Diagram of workspace setup and target locations for retrieval strategies in Experiment 2. The feedback type and workspace setup remain consistent with the algorithmic group. But participants were required to aim directly at the Non-Critical target location. C) Hand angle for the Critical target locations. D) Absolute hand angle error for the Non-Critical target location. E) Reaction time for the Critical target location. F) Reaction time for the Non-Critical target location. G) Generalization curves for algorithmic and retrieval strategies. Beneath the curves, bars represent the distribution of reaches for each group. The shaded gray region marks the “aiming zone,” where participants were most likely to aim when counteracting the 45° rotation at the Critical target. The amplitude, mean, and width parameters displayed in the figure were derived from Gaussian function fits.

#### Adaptation performance

Because the feedback type was manipulated at all Non-Critical targets, we first asked whether this manipulation changed overall performance in either group. As in Experiment 1, participants in Experiment 2 adapted rapidly when the visuomotor rotation was introduced, and hand angles remained largely stable from early to late training (Figure 3C, Figure S2A for trial-by-trial hand angles time courses). Note, caching of the strategy was facilitated by the 5-trial pre-exposure to the 45° visuomotor rotation for the retrieval group; this was intentionally implemented to minimize any potential influence of a fleeting algorithmic strategy during the early adaptation period. Using the same analysis as Experiment 1 to compare learning functions, we found an overall group difference: the retrieval group produced smaller hand angles than the algorithmic group (main effect of Group: b = 2.46, SE = 0.92, t(76) = 2.66, p = .001). This effect was driven primarily by early training. The algorithmic group initially overshot the required compensation (50.18 ± 9.43°), producing larger hand angles than the retrieval group (44.84 ± 6.98°; p = .02). By late training, the groups converged, with both showing slight undershoot relative to the imposed 45° rotation (algorithmic: 46.58 ± 8.96°; retrieval: 42.10 ± 3.16°), and the between-group difference was comparable (p = .06). As expected, we did observe clear evidence that adaptation changed over time (main effect of Time: b = 1.58, SE = 0.71, t(76) = 2.24, p = 0.03), but we did not observe Group x Time interaction: b = 0.22, SE = 0.71, t(76) = 0.31, p = .76. Furthermore, the stability of a cached memory was more stable or precise than executing an algorithmic strategy, as reflected by a main effect of Group (variability for retrieval - algorithmic: b = 1.80, SE = 0.74, t(76) = 2.44, p = .002).

Performance on the Non-Critical targets was essential to ensure that the strategies used to counteract the imposed rotation at the Critical target location in each group were dissociated. Because participants in the algorithmic group were required to compensate for the 45° rotation at the Non-Critical targets, whereas participants in the retrieval group were instructed to aim directly at those target locations, we quantified absolute error relative to each target location so that performance could be compared numerically across groups. At the Non-Critical target locations, the analysis revealed a significant main effect of Group (b = 1.83, SE = 0.78, t(76) = 2.33, p = .02) and a significant Group × Time interaction (b = 1.03, SE = 0.48, t(76) = 2.12, p = .04), but no main effect of Time (b = 0.50, SE = 0.48, t(76) = 1.04, p = .30). Planed linear contrast analysis indicated that absolute error decreased from early to late training in the algorithmic group (9.10 ± 10.08° vs. 6.04 ± 4.76°, p = .03), whereas it remained stable in the retrieval group (3.39 ± 3.13° vs. 4.44 ± 3.07°, p = .45). Accordingly, the retrieval group showed more better performance than the algorithmic group during early training (p < .01), but this group difference was no longer present by late training (p = .39) (Figure 3D; Figure S2B for the complete hand angles time course). Together, these results confirm that task performance in both groups was consistent with the instructed strategy requirements.

#### Preparation time

As both groups presented comparable learning functions after a 45° rotation, it was crucial to verify whether distinct strategies were adopted within each group by comparing their RT profiles. Compared to the average baseline RT, both the algorithmic (p < 0.01) and retrieval groups (p < 0.01, Figure S2E-F for RT comparison between baseline and training) showed increased RT during the training. Consistent with Experiment 1, both groups reached a comparable level of performance, but did so with different computational costs, as reflected in reaction time (Figure 3E, Figure S2C). Indeed, the main effect of Group indicated that the retrieval group responded faster than the algorithmic group across training (b = 0.21, SE = 0.04, t(76) = 5.03, p < .001). There was also a significant main effect of Time (b = 0.15, SE = 0.03, t(76) = 5.04, p < .001), indicating that reaction times decreased from early to late practice. More interestingly, the analysis detected a significant Group × Time interaction (b = 0.06, SE = 0.03, t (76) = 2.23, p = 0.03). This overall speeding was driven mainly by the algorithmic group, whose responses became markedly faster with training (early: 1.15 ± 0.54 s; late: 0.88 ± 0.27 s; p < .001). In contrast, the retrieval group showed modest change over time, with reaction times remaining more stable from early to late practice (0.56 ± 0.15 s to 0.49 ± 0.12 s; p = .05) than that in the algorithmic group. This pattern of results indicated that the behavioral signatures at the Critical target location were consistent with those observed in Experiment 1 despite the stricter controls.

Further, we validated reaction times at the Non-Critical targets as an additional check that the revised design continued to elicit distinct strategies in the two groups. As expected, we found a significant main effect of Group (b = 0.24, SE = 0.04, t (76) = 5.80, p < .001), indicating that the algorithmic group had greater RTs than the retrieval group on the Non-Critical targets. The significant main effect of Time (b = 0.09, SE = 0.03, t (76) = 3.40, p < .001) suggested a trend toward reduced RT over practice. The main effect of the Group × Time interaction approached statistical significance (b = 0.05, SE = 0.03, t (76) = 1.98, p = .051; Figure 3F, Figure S2D), indicating that the change in RT across groups was comparable. These results show that the RT signatures were consistent across target locations, reinforcing that our design manipulation successfully induced distinct strategies in the two groups.

#### Implicit recalibration

The primary goal of Experiment 2 was to test whether different forms of explicit strategy differentially shape implicit recalibration at the Critical target location when implicit recalibration at all Non-Critical target locations was minimized to prevent possible error spillover. Compared with Experiment 1, Experiment 2 included two additional modifications to strengthen experimental control: (1) the angular separation between the Critical target and the adjacent Non-Critical targets was increased from 60° to 90°, and (2) delayed endpoint feedback was applied at all Non-Critical targets to suppress implicit recalibration at those locations. With these additional controls in place, we sought to isolate the unique influence of strategy on implicit recalibration at the Critical target.

We first compared implicit recalibration at the Critical target probe location between the algorithmic and retrieval groups to test whether strategy use directly modulates its magnitude. Numerically, implicit recalibration at the Critical target was greater in the retrieval group (15.49 ± 8.99°) than in the algorithmic group (11.43 ± 6.43°), but this difference did not reach significance, t(38) = −1.65, p = .11 (Figure 3G). Consistent with this result, Gaussian fits to the generalization functions revealed comparable amplitudes across groups (algorithmic: 14.99°; retrieval: 16.20°; bootstrap, p = .28; Figures 3G; see Figure S5B for model fitting). In contrast, the two groups exhibited distinct centers of generalization (algorithmic: µ = 28.69°; retrieval: µ = 10.94°; bootstrap, p = .01, Figure S5B), a difference that likely reflects their markedly different distributions of aiming directions and is consistent with the principle of plan-based generalization (McDougle et al., 2017).

Importantly, the stricter controls introduced in Experiment 2 reduced the breadth of the algorithmic group’s generalization function (σ = 53.06°) relative to Experiment 1 (σ = 75.41°). This narrowing is consistent with our feedback manipulation successfully constraining implicit recalibration at the Non-Critical target locations, as evidenced by comparable aftereffects at those locations in the two groups (algorithmic: 5.66 ± 4.31°; retrieval: 4.05 ± 3.58°), t (38) = 1.28, p = .21 (Figure S2G-H). Notably, the aftereffect of the Non-Critical targets (5.66 ± 4.31°) in the algorithmic group was reduced significantly compared to the algorithmic group in Experiment 1 (15.00 ± 8.36°). Moreover, relative to the aftereffect measured at the Critical target, aftereffects at the Non-Critical target locations were significantly reduced to negligible levels in both the algorithmic and retrieval groups (both p < .01; Figure S2G-H). By contrast, the breadth of the retrieval group’s generalization function (σ = 37.12°) remained similar to that observed in Experiment 1 (σ = 32.97°), consistent with the fact that participants in this group simply reached directly toward the assigned target locations. Together, the selective narrowing observed in the algorithmic group provides strong evidence that the statistics of planned aiming directions can reshape implicit recalibration through plan-based generalization.

Nevertheless, the difference in the breadth of the generalization function between the algorithmic and retrieval groups remained significant (Figures 3G; bootstrap, p = .02), with the algorithmic group exhibiting a broader generalization function than the retrieval group. However, this difference appeared to be markedly reduced relative to that observed in Experiment 1 (Figure S5B). First, this modest difference in generalization breadth cannot be explained by overall adaptation level: adaptation was comparable between groups immediately before entering each Exclusion block (F (1, 256) = 3.04, p = 0.08) and did not change reliably across trials (F (6, 256) = 1.14, p = .34, Figure S4B for comparison at the “top-up” trials). Second, the group difference in generalization is unlikely to reflect continued use of algorithmic strategies during the Exclusion blocks. Comparing RTs in the Exclusion block across groups, the analysis showed no main effects of Group (F (1, 266) = 3.15, p = 0.08) or Time (F (6, 266) = 1.04, p = 0.40, Figure S2C-D, Figure S4E for comparison of Exclusion RTs), indicating that both groups responded comparably quickly. To further verify that participants withheld the strategy during the Exclusion block, we compared RTs during Exclusion with baseline RTs and training RTs within each group. In the algorithmic group, RT during the Exclusion block (0.50 ± 0.13 s) was significantly longer than at baseline (0.43 ± 0.14 s, p = .03), but markedly shorter than during training (1.02 ± 0.34 s, p < .01, Figure S2E-F). In the retrieval group, RT during the Exclusion block (0.43 ± 0.10 s) was comparable to baseline RT (0.40 ± 0.15 s, p = .62), yet significantly shorter than RT during training (0.50 ± 0.13 s, p < .01, Figure S2E-F). This pattern suggests that participants attempted to withhold the strategies used during training during the Exclusion block, as evidenced by the reduction in RT relative to training in both groups.

While a number of potential confounds were controlled for in Experiment 2, there nonetheless a lingering difference in the variance of distribution associated with reaches at the Critical target location, as the algorithmic group exhibited greater variability than the retrieval group (Figures 3G; bootstrap, p < .001). Consequently, stricter control of plan-based generalization from the Non-Critical targets, achieved by restricting online feedback, yielded broadly comparable generalization functions across the algorithmic and retrieval groups. However, these constraints were insufficient to equalize the variance of the distribution of reaches at the Critical target, which likely contributed to the remaining modest group difference (Figure 3G, Figure S5B).

### Experiment 3: Under tight control of sensory prediction errors, do differences in implicit recalibration vanish?

In Experiments 1 and 2, we found that variability in reach directions strongly influenced the breadth of implicit generalization. Consistent with the plan-based generalization account, implicit recalibration is expected to peak around the planned aiming direction (McDougle et al., 2017). Greater variability in reach direction likely reflects a broader distribution of planned aiming directions, given that the total reach direction reflects the combined contributions of explicit re-aiming and implicit recalibration (Taylor and Ivry, 2014; McDougle et al., 2015), whereas the implicit learning function itself is thought to remain relatively stable. This creates a critical interpretive challenge: when variance changes, the generalization profile can change with it, making it hard to isolate what each explicit strategy contributes. To provide the strictest control over reach variance and error feedback that could spill over to the implicit recalibration system, in Experiment 3 we leveraged an error-clamp paradigm to generate sensory prediction errors that were incidental to task goals (Morehead et al., 2017). As the error-clamp paradigm renders explicit re-aiming strategies irrelevant, we can gain complete control over the reach variance along with the sensory-prediction errors that drive implicit recalibration.

In Experiment 3, participants reached to nine different locations in the workspace, defined as angular locations ranging from 45° to 165° relative to the Critical target, which was always positioned at the top of the workspace (Figure 4A). This design was intended to preserve the workspace configuration used in Experiments 1 and 2 while incorporating the error-clamp paradigm into the same spatial context. To dissociate strategy use, we manipulated instructions for reaching and, importantly, the visibility of the target locations between the groups during training. For the algorithmic group, all reach locations except for the Critical target were invisible, requiring participants to rely entirely on mental rotation to compute the instructed reach direction relative to the Critical target. The intended reach direction was provided via text-based instructions. For example, if participants saw “move towards 45°”, they were expected to attempt to reach 45° away from the critical target but they did not see a visual target location at 45° (Figure 4A). Similarly, if participants saw “move towards 75°” (Figure 4B) or “move towards 105°” (Figure 4C), they were expected to reach toward 75° or 105°, respectively, without any visual target. Importantly, when participants were instructed to reach toward 45°, their reach was accompanied by 15° error-clamped cursor feedback (Figure 4A); reaches toward all other instructed locations received no cursor feedback (Figure 4B – C). For all these instructed and invisible reach locations, the accuracy of reach performance was indicated by computer-generated verbal feedback with 4 different scales (“Excellent”: absolute error < 5; “Good move”: 5< absolute error <10; “Fair move”: 10 < absolute error < 15; “Poor move”: absolute error > 15).

**Figure 4.**
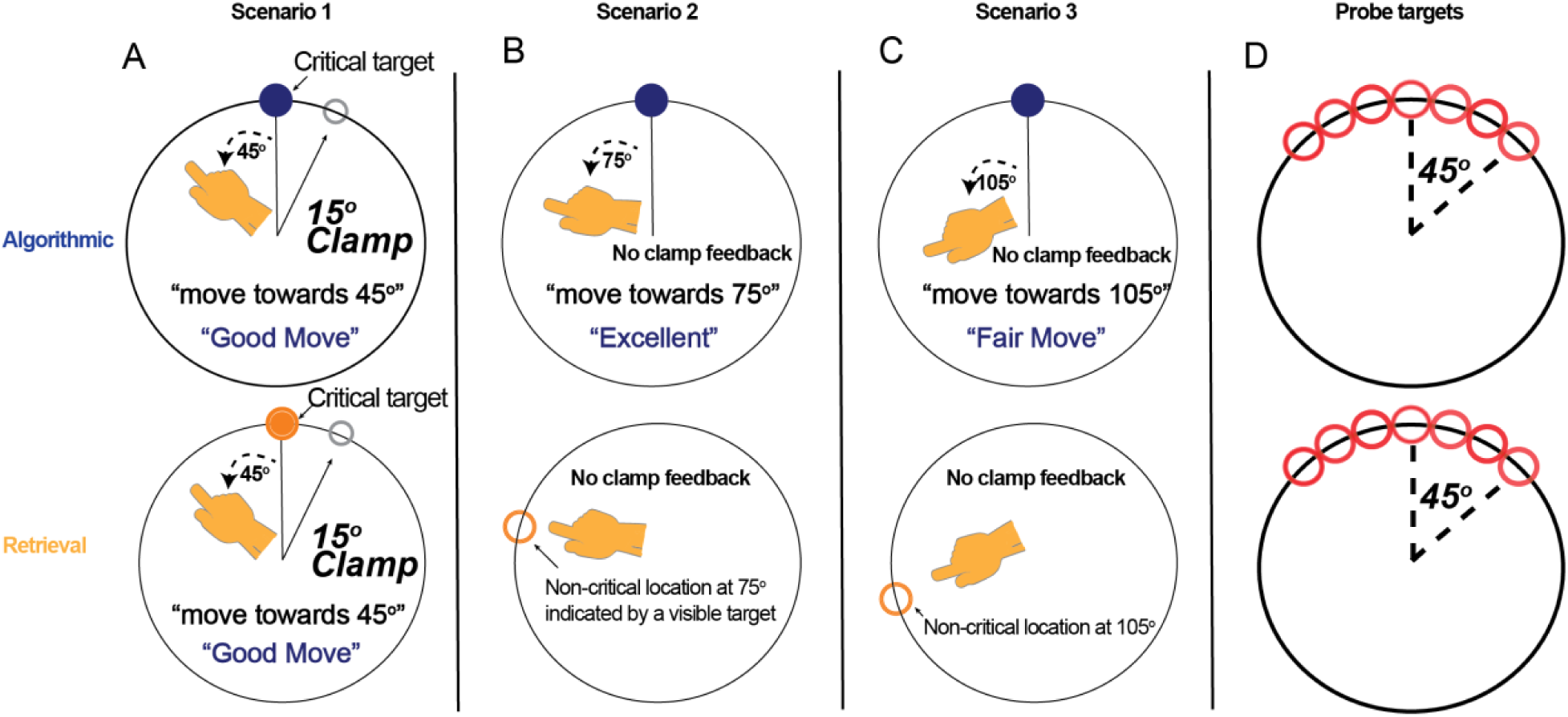
Experimental design incorporating an error-clamp paradigm to assess the influence of strategy use on implicit recalibration. The examples of particular workspace scenarios used in Experiment 3 for the algorithmic and retrieval groups. A) The scenario for trials reaching toward the 45° location relative to the Critical target with 15° error-clamped feedback. Note that both groups lack a landmark for the 45° location, and only this scenario has the error-clamped feedback. B) The scenario for trials reaching toward the 75° relative to the Critical target. For the algorithmic group, the 75° location relative to the Critical target must be computed using a mental rotation strategy because no visual target is presented. In contrast, for the retrieval group, a target is displayed at the location that is 75° relative to the Critical target. Participants were asked to simply reach toward it. C) The scenario for trials requiring reaching toward the 105°; no target is present for the algorithmic group while a target is presented for the retrieval group. For any Non-Critical angular direction (any reach location other than 45°), no cursor feedback or clamped feedback was imposed. D) The 7 probe targets (including the Critical target) are selected for measuring implicit recalibration during the Exclusion blocks.

For the retrieval group, when the Critical target was presented, participants were instructed to “move toward 45°” and their reach was accompanied by 15° error-clamped cursor feedback, which matches the instruction and feedback for the Critical target for the algorithmic group (Figure 4A). This enabled implicit recalibration to operate on the error clamped feedback while planning a reach towards 45° relative to the Critical target. For all other instructed reach locations the targets were visible, allowing participants to refrain from any strategy use or mental rotation for these Non-Critical reach locations. Importantly, because reaching toward these Non-Critical locations did not include error-clamp feedback, this design eliminated the possibility that implicit recalibration at the Critical target could result from error spillover from planning reaches at the Non-Critical locations (Figure 4B, C). This design was intended to parallel the algorithmic and retrieval conditions of Experiments 1 and 2, while affording full control over the reach distribution (via instructions) and sensory-prediction errors. The goal of this design was to minimize the extent to which the resulting generalization function could be shaped by movement dispersion, thereby providing a more direct test of how algorithmic and retrieval strategies differentially shape implicit recalibration. Implicit recalibration for both the algorithmic and retrieval groups at probe target neighboring the Critical target was measured during Exclusion trials where participants were instructed to reach directly to the target without any cursor feedback (Figure 4D).

#### Adaptation performance

When reaching to the 45° location, hand angles were comparable across groups, with no main effect of Group (b = -0.30, SE = 0.76, t (72) = −0.39, p = .70). Participants did show a modest decline in hand angle from early to late practice (b = 1.31, SE = 0.71, t (72) = 1.84, p = .07), but this implicit drift was similar in both groups (group × time: b = 0.90, SE = 0.71, t (72) = 1.27, p = .21). In other words, despite the design differences from all previous experiments, the algorithmic (43.07 ± 4.99°) and retrieval groups (45.47 ± 3.69°) ultimately reached an essentially comparable level of adaptation at the 45° location in the end of training (Figure 5A, Figure S3A). Furthermore, a cached memory from the retrieval group (6.25 ± 1.73°) tended to be more stable than executing an algorithmic strategy (7.86 ± 3.25°), as reflected by a main effect of Group (b = 0.81, SE = 0.41, t(72) = 1.96, p = .05).

**Figure 5.**
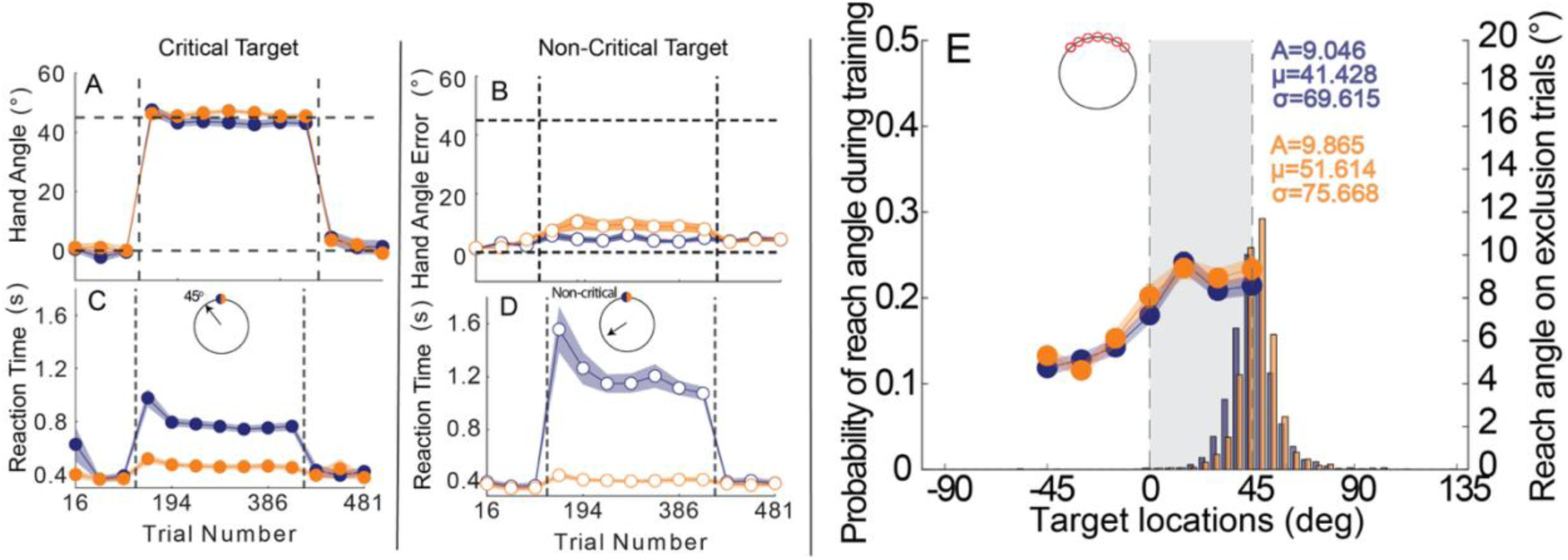
With the strongest control for confounds, implicit generalization function becomes comparable between two groups. A) Hand angles at the 45° location relative to the Critical target. B) Absolute hand angle errors at the Non-Critical locations. C) Reaction times at the 45° location relative to the Critical target. D) Reaction time at the Non-Critical locations. E) Generalization curves for algorithmic and retrieval strategies. Beneath the curves, bars represent the distribution of reach direction for each group. The shaded gray region marks the “aiming zone,” where participants were most likely to aim when reaching the 45° target relative to the Critical target. The amplitude, mean, and width parameters displayed in the figure were derived from Gaussian function fits.

To provide an additional check that performance was matched across groups under error-clamp conditions, we also examined absolute errors when reaching the Non-Critical locations relative to the Critical target across early and late practice (Figure 4B – C). The absolute error can provide a numerically equivalent measure for comparing performance between the algorithmic and retrieval groups, as they performed relatively distinct tasks under different conditions. The algorithmic and retrieval groups performed similarly (no main effect of Group: b = -1.17, SE = 0.65, t(72) = -1.81, p = 0.07), and errors remained stable over time (no main effect of Time: b = 0.10, SE = 0.58, t(72) = 0.18, p = 0.86), with no group differences in change across practice (Group × Time: b = 0.33, SE = 0.58, t(72) = 0.58, p = .56) (Figure 5B, Figure S3B). Therefore, both groups accomplished the task with comparable performance in reaching assigned locations relative to the Critical target.

#### Preparation time

Despite the major redesign in Experiment 3, the reaction-time pattern at the 45° Critical location closely matched what we observed in earlier experiments while adapting 45° rotation at the Critical target (Figure 5C). Across groups, responses became faster from early to late training, yielding a reliable main effect of Time (b = 0.07, SE = 0.01, t(72) = 5.58, p < .001). We also observed a strong main effect of Group (b = 0.19, SE = 0.03, t(72) = 7.54, p < .001), indicating that the retrieval group responded markedly faster than the algorithmic group overall. At the same time, the magnitude of improvement differed between groups, as reflected by a significant Group × Time interaction (b = 0.37, SE = 0.01, t(72) = 2.98, p = .004). This interaction was driven primarily by a pronounced reduction in reaction time for the algorithmic group from early (0.98 ± 0.29 s) to late training (0.76 ± 0.14 s, p < .001), whereas reaction time in the retrieval group changed little (early: 0.52 ± 0.13 s; late: 0.45 ± 0.09 s, p = .07). Notably, this dissociation persisted even without a visible target at the 45° location relative to the Critical target. The retrieval group’s consistently fast responses are consistent with caching a stable aiming solution, whereas the algorithmic group appeared to rely on trial-by-trial computation of the appropriate aiming direction. In addition, RTs in the algorithmic group were significantly faster during training than at baseline (p < .01, Figure S3E), whereas RTs in the retrieval group were comparable between training and baseline (p = .07, Figure S3F), consistent with efficient retrieval of the correct aiming strategy.

Further, we validate this interpretation that the two groups relied on distinct explicit strategies to plan their aiming solutions by comparing their RT profiles while reaching the Non-Critical locations relative to the Critical target (Figure 5D, Figure S3D for the complete hand angle time course). The RT profile largely mirrored what we observed when aiming at the 45° location. In particular, both groups increased their response speed from early to late training (main effect of Time: b = 0.13, SE = 0.03, t(72) = 3.39, p < 0.01), with the retrieval group having an overall faster RT than the algorithmic group (main effect of Group: b = 0.44, SE = 0.05, t(72) = 8.25, p < .001). In addition, the amount of improvement differed by Group (Group × Time: b = 0.11, SE = 0.03, t(72) = 3.39, p < 0.01): the algorithmic group showed a clear reduction in RT across practice (early: 1.56 ± 0.75 s; late: 1.08 ± 0.21 s; p < .001), whereas the retrieval group remained consistently fast with little change (early: 0.46 ± 0.10 s; late: 0.42 ± 0.08 s; p = 0.73). This replication, while reaching Non-Critical locations, provides converging evidence that Experiment 3 successfully dissociated the explicit strategies used by the two groups.

#### Implicit recalibration

As in the first two experiments, the primary goal was to determine whether different kinds of explicit strategies have downstream effects on implicit recalibration. We first examined whether different kinds of strategies differentiate their impact on the amplitude of implicit recalibration at the Critical target location. As a result, we found there was no significant difference in amplitude of implicit recalibration arising from different kinds of explicit strategies (Figure 5E; algorithmic: 7.21± 3.36°, retrieval: 8.08 ± 4.24°, t(36) = -0.70, p = 0.49). This result was further supported by fitting hand angles from Exclusion trials to a Gaussian function (Figure S5C for model fitting), indicating that the amplitude between algorithmic (A = 9.05 °) and retrieval groups (A = 9.87 °) was comparable (Figure S5C). Likewise, the center of the generalization function was statistically indistinguishable between groups (Figures 5E; algorithmic: µ = 41.43°, retrieval: µ = 51.61°; bootstrap, p = 0.37). Notably, under the error-clamp design, the breadth of generalization also converged: the algorithmic group (σ = 69.62°) and retrieval group (σ = 75.67°) showed comparable breadths (Figures 5E; bootstrap, p = 0.39). While the algorithmic group continued to generate more variable aiming solutions than the retrieval group (Figure S6C, bootstrap, p < 0.01), they did not receive any accompanying error-based cursor feedback that could train the implicit recalibration system.

Again, this comparison of generalization function was based upon the fact that both groups reach comparable levels of adaptation prior to Exclusion trials (F (1, 243) = 3.26, p = 0.07), which was sustained throughout the entire training period, reflected by a main effect of Trials (F (6, 243) = 0.74, p = .62, Figure S3A – B, Figure S4C). In addition, RTs between the algorithmic and retrieval groups were comparable during the Exclusion block measuring implicit recalibration (F(1, 252) = 0.13, p = 0.72). Relative to training RTs, RTs during the Exclusion block were significantly faster in both the algorithmic (p < .01, Figure S3E) and retrieval groups (p < .01, Figure S3F), suggesting that participants attempted to withhold the strategies engaged during training. Relative to the 45° location, the aftereffect measured while reaching Non-Critical location was substantially reduced in both the algorithmic (normalized to baseline; 45°: 5.24 ± 8.26°, Non-Critical: 0.65 ± 3.17°; p = 0.01; Figure S3G) and retrieval groups (normalized to baseline; 45°: 5.50 ± 8.86°, Non-Critical: 0.96 ± 4.00°; p = 0.03; Figure S3H), suggesting little implicit recalibration occurred at all Non-Critical locations. These aftereffects measured at the 45° location were significantly greater than baseline in each group (all ps = 0.01).

#### The variability of reaches, not strategy use, reshapes the implicit landscape

In Experiments 1 and 2, we found broader generalization of implicit recalibration when participants used an algorithmic strategy compared to a retrieval strategy, which echoes the broader generalization observed for algorithmic strategies in relative isolation from implicit recalibration (McDougle and Taylor 2019). However, we found that when participants used an algorithmic strategy, they also had a greater distribution of reaches in the workspace. Because these reaches were accompanied by cursor feedback it could result in error spillover to the implicit recalibration system. In Experiment 3, when we strictly controlled both the reach distribution and error feedback, we found that the generalization function of implicit recalibration was the same regardless of the kind of strategy use. To further characterize the relationship between generalization and reach variance, we examined the residual of these functions between the retrieval and algorithmic conditions across experiments. (Figure 6). We found that the residual of the generalization function (Figure 6A) mirrored the residual of the reach distributions (Figure 6B). Indeed, as the experimental task design exerted increasingly more control over the reach and error distribution, the differences in the width (Figure 6C) of the fitted generalization function decreased. The center (Figure 6D) and amplitude (Figure 6E) of the generalization of implicit recalibration always remained the largely unchanged. This direct comparison supports the interpretation that the broader implicit generalization function observed with the algorithmic strategy arises from error spillover, not from the kind of strategy use, which is consistent with predictions of plan-based generalization (Day et al., 2016; McDougle et al., 2017).

**Figure 6.**
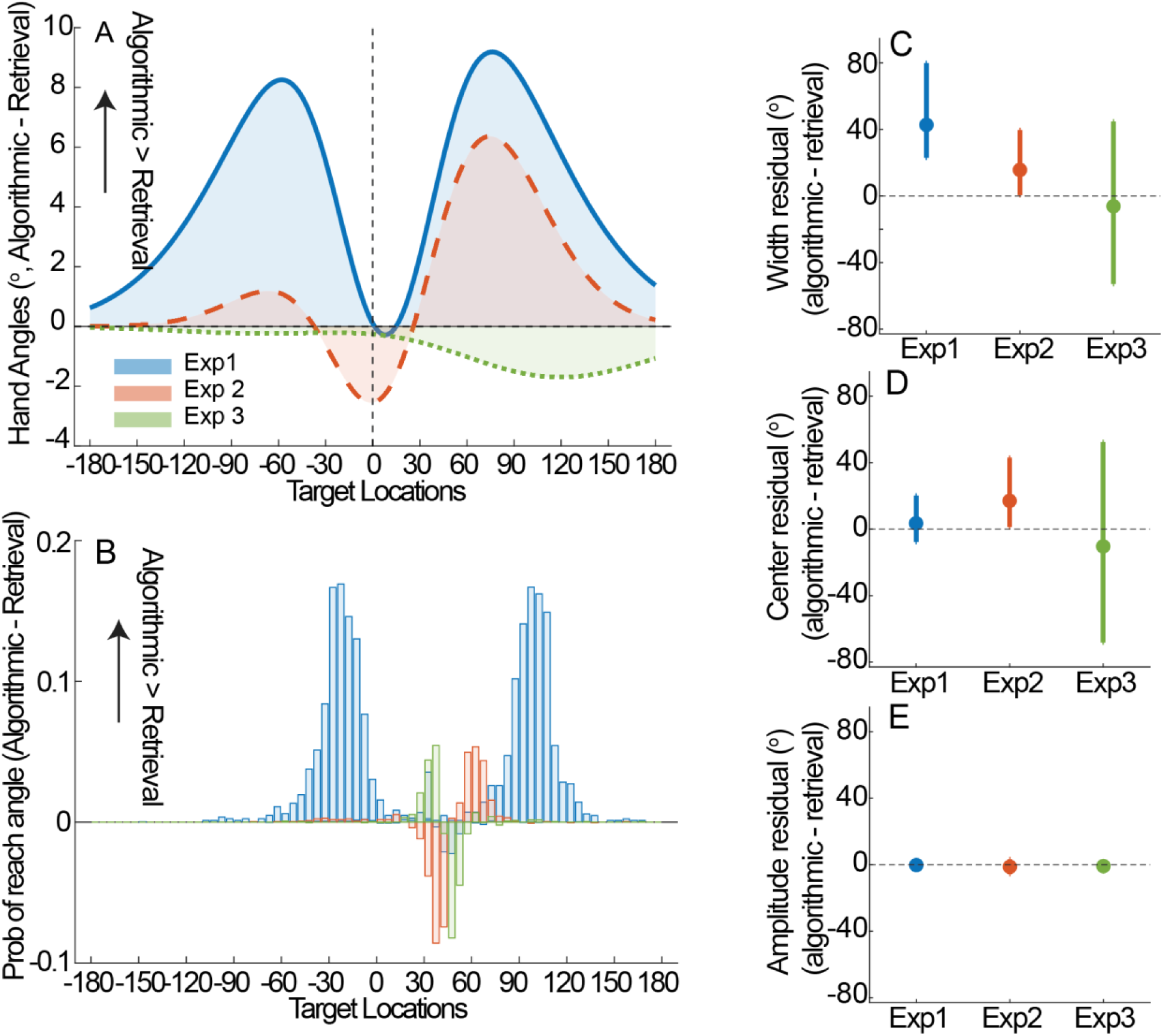
Residual analysis of the implicit generalization function by subtracting retrieval from algorithmic. A) Contrast of generalization function after fitting the model using algorithmic - retrieval strategies. X axis represents probing target locations, and Y axis represents the magnitude of hand angle. The positive value indicates that algorithmic strategy resulted in greater implicit recalibration at the probe target location. B) The difference in probability of reach direction across workspace (−180° – 180°). A positive value indicates that algorithmic strategies reach that area more frequently. In Exp1, there were no implicit recalibrations at the Non-Critical target in the retrieval group. When comparing the probability of reach directions for the Non-Critical target, we used the Algorithmic strategy minus zero. The implicit recalibration at the Non-Critical targets showed a clear spillover effect because the residual of reach-angle probability is highly consistent with the residual of generalization function. C) Residual width of the generalization function across all three experiments. From Experiment 1 to 3, the width of generalization function gradually reduced with the extent of experimental control. D) Residual center of the generalization function across all three experiments. E) Residual amplitude of the generalization function across all three experiments. In C – E, data was derived from bootstrap, and error bars indicate the 95% confidence interval.

## Discussion

### Summary

In this study, we tested whether algorithmic and retrieval-based strategies, both of which are known to improve performance in visuomotor adaptation, differentially affect implicit recalibration – a central debate in the field. Given that these two strategies likely rely on distinct neural circuits, they could interact with the cerebellar-based recalibration system in different ways. Furthermore, retrieval strategies form stricter stimulus-response associations compared to algorithmic strategies, which could reshape implicit recalibration at a more process level. Indeed, a prior study has found greater generalization of the strategy under algorithmic compared to retrieval strategies but the study deliberately minimized implicit recalibration, leaving open the question of whether such strategy differences truly penetrate implicit recalibration (McDougle and Taylor, 2019).

To address this question, we manipulated the target set size to bias participants towards algorithmic and retrieval strategies (McDougle and Taylor, 2019; Velazquez-Vargas and Taylor, 2024), while controlling for several confounding factors, such as inter-trial interval and memory decay (Hadjiosif et al., 2023), plan-based generalization (Day et al., 2016; McDougle et al., 2017; Chen and Taylor, 2025), and unequal training exposure (Bond and Taylor, 2015). To quantify implicit recalibration, we interleaved periodic Exclusion trials in which error feedback was withheld, and participants were instructed to withhold any explicit strategy (Werner et al., 2015; Maresch et al., 2021; t Hart et al., 2024; Chen and Taylor, 2025). Our experimental design across a set of experiments was successful in dissociating the type of strategy used, with reliably slower RTs in the algorithmic group, yet both groups reached comparable asymptotic adaptation (∼45°). These important experimental controls were essential in supporting that any observed differences in implicit recalibration reflected only differences in strategic demands rather than differences in exposure or target statistics.

In Experiments 1 and 2, we observed a broader implicit generalization function under algorithmic than retrieval strategies. Although this pattern could be construed as consistent with classic accounts in which stimulus variability promotes generalization (Estes and Burke, 1953; Schmidt, 1975), we do not think it reflects a fundamental change in implicit recalibration per se. Instead, we interpret the apparent broadening as a consequence of plan-based generalization: because algorithmic strategies generate more variable movement plans, implicit recalibration may accrue across variable aiming locations, producing error spillover that inflates the measured breadth of the implicit generalization function around the Critical target. Critically, in Experiment 2, we imposed stricter constraints to minimize implicit recalibration at Non-Critical targets. This manipulation substantially reduced the distribution of learning-relevant errors (training inputs) across the workspace, and the measured implicit generalization function narrowed accordingly. Because implicit recalibration has been shown to generalize around the planned movement (Day et al., 2016; McDougle and Taylor, 2019; Poh and Taylor, 2019), any residual broadening under algorithmic strategies would be expected if participants continued to produce more variable aiming solutions at the Critical target. Consistent with this account, our complementary analyses of reaching distribution showed that hand angles at the Critical target were more dispersed under algorithmic than retrieval strategies. This pattern supports the perspective that explicit strategy functions as a controller that shapes the training inputs received by the implicit system (Schween et al., 2018; Poh et al., 2021). Once variability characterized by explicit strategies becomes irrelevant (Experiment 3), the difference in generalization function becomes negligible. Modern probabilistic accounts similarly argue that generalization reflects inference about error sources and the relevance of disturbances, again emphasizing statistics of experience rather than the subjective strategy used to generate actions (Berniker and Kording, 2008). Our results support this account: the use of algorithmic and retrieval strategies can produce differing spatial distributions of these re-aiming plans and, as a consequence, implicit generalization follows those plan statistics. When we removed such a feedthrough pathway with error clamp, the implicit map no longer differed across groups (Figure 4G). Again, our results reinforce the view that explicit and implicit learning operate relatively independently, besides the indirect effects of being in tandem with one another (Taylor and Ivry, 2011; Miyamoto et al., 2020). On this account, explicit strategies need not exert any “special” influence over implicit recalibration beyond their role in shaping the state the learner occupies.

### The interaction of explicit and implicit adaptation

The literature offers compelling, yet mixed, accounts of how explicit and implicit processes interact. One prominent view is that explicit re-aiming strategies and implicit recalibration reflect largely independent learning processes, such that overall adaptation can be approximated as the sum of these components (Mazzoni and Krakauer, 2006; Taylor and Ivry, 2011, 2014). Support for this account comes from aiming-report and error-clamp paradigms, which suggest that implicit recalibration often unfolds in a relatively stereotyped manner and shows limited sensitivity to factors such as task relevance and error size (Bond and Taylor, 2015; Morehead et al., 2017; Kim et al., 2018). Under this view, the explicit component acts as a flexible compensatory controller, adjusting in response to any residual task performance error (Taylor and Ivry, 2011). Implicit recalibration can also respond to errors produced by explicit re-aiming (Miyamoto et al., 2020). Furthermore, implicit recalibration appears to be inextricably yoked to where explicit re-aiming strategies operate – referred to as plan-based generalization (Day et al., 2016; McDougle et al., 2017). These findings suggest that potential interactions between explicit strategies and implicit recalibration are simply indirect because they are operating in series (Taylor and Ivry, 2011; McDougle et al., 2017; Miyamoto et al., 2020).

An alternative view puts forth that explicit and implicit processes compete for a common error signal, where, reducing one component is often accompanied by a compensatory increase in the other, yielding an inverse relationship between explicit and implicit contributions (Albert et al., 2022). This pattern is typically interpreted as evidence for a direct interaction – rather than independence – between the two processes, which can lead to more complex or, at least, nonlinear interactions when explicit strategies are at play in visuomotor rotations tasks (Neville and Cressman, 2018; t Hart et al., 2024) The nature of this potential interaction is especially complicated because it depends on how these components are operationalized and measured during adaptation. Multiple studies argue that common assessment methods can bias or distort estimates of explicit and implicit contributions, complicating inferences about their interaction (Maresch et al., 2021; t Hart et al., 2024). At present, we do not have a way to probe explicit strategies or implicit recalibration that does not result in some sort of an intervention or observer effect. Even leveraging approaches, such as limiting preparation time (Haith et al., 2015), do not offer a way to independently measure explicit and implicit learning without breaking the fourth wall (Chen and Taylor, 2025).

Here, we attempted to, at least avoid, these potential intervention problems by first letting training to unfold naturally under our different task demands before introducing the Exclusion trials to probe implicit recalibration. Target set size alone was sufficient in biasing participants towards different strategies, as measured indirectly from RT. We then implemented a stepwise set of increasingly stringent controls across Experiments 1–3 and ultimately found that algorithmic and retrieval strategies produced progressively indistinguishable functions of implicit recalibration. Under error-clamp conditions – where implicit updating is largely insulated from plan-based generalization – implicit recalibration showed little sensitivity to the underlying form of the strategy, consistent with an independent account of explicit and implicit processes in sensorimotor adaptation (Mazzoni and Krakauer, 2006; Taylor and Ivry, 2011; Taylor et al., 2014; Morehead et al., 2017).

Beyond directly affecting implicit recalibration, a number of studies have demonstrated alternative routes through which explicit strategies could act as contextual cues to create, select, and update a different motor memory (Heald et al., 2021). In particular, the motor system appears to segment experience into latent causes and uses available cues to infer which memory should be expressed and updated; a strategy could therefore become part of the cue set only if it shifts the inferred latent structure of the task (Heald et al., 2021). For example, the study showed broader generalization when targets were rated as highly similar, suggesting that explicit strategy selection effectively treated them as the same context (Poh et al., 2021). Moreover, when participants were encouraged to prioritize one contextual cue over another, generalization increased specifically along the dimension that was emphasized (Poh et al., 2021). In our experiments, participants always faced the same overarching aiming problem under a stable task framing, and the key manipulation was whether the solution was computed online or retrieved from memory. That distinction may not have constituted a salient contextual boundary for implicit learning to assign error to distinct latent causes (Heald et al., 2021). However, if different strategies were embedded within meaningfully different inferred task states, such as distinct goals, rules, mappings, or conceptually separable contexts, then strategy adoption could become a strong contextual signal, potentially supporting more separable implicit memories even when movement plans overlap (Heald et al., 2021; Poh et al., 2021). This possibility is especially relevant for paradigms in which implicit adaptation contributes a larger fraction of behavior (e.g., dynamic force-field adaptation; (Forano et al., 2021)). If the form of strategy can serve as a contextually-bound cue to retrieve the appropriate implicit mapping, then the degree to which strategies directly or indirectly affect implicit recalibration becomes more of a categorical or philosophical distinction.

## Conclusion

By rigorously controlling multiple confounds that influence implicit recalibration across three experiments, our findings suggest that the strategies employed during visuomotor adaptation do not differentially reshape the implicit recalibration landscape. Explicit strategies do not appear to directly impact implicit recalibration beyond the observation that they simply act as a gate to control the data inflow that is used to update the implicit mapping. Simply put, the implicit recalibration system will faithfully update its mapping with whatever training data it is provided. Our findings offer a more pedestrian explanation for the role of training variability or, more broadly, guidance theories for generalization (Schmidt, 1975) – it is just the sampling distribution when it pertains to implicit sensorimotor adaptation.

## Material and Methods

### Participants

A total of 164 participants (66 males, 98 females; mean age = 24.66 years, SD = 3.79) were recruited through the Princeton University psychology subject pool in exchange for course credit or from the local community in exchange for monetary reimbursement for their time. All participants were right-handed, confirmed by the Edinburgh Handedness Test (Oldfield, 1971), and reported normal or corrected-to-normal vision. The study protocol was approved by the Institutional Review Board at Princeton University, and all participants provided written informed consent prior to participation.

### Task and Apparatus

Participants performed a center-out visuomotor reaching task using a stylus on a Wacom digitizing tablet. Their goal on each trial was to bring a virtual cursor to a visually displayed target by sliding the stylus across the tablet surface. The cursor and targets were presented on a 17 inch Dell LCD monitor (1024 × 768 resolution; 60 Hz refresh rate) running Windows 7. The monitor was positioned 10 inches above the tablet, fully occluding direct vision of the hand to ensure that participants relied solely on the displayed cursor for visual feedback (Figure 1A). Each trial began with the participant being guided to a central starting location (5-mm diameter). A white ring dynamically expanded or contracted with the radial distance of the hand from the center to cue participants toward the start position without precise angular cursor feedback. When the hand came within 6 mm of the center, a white cursor (5 mm in diameter) appeared to more precisely guide the hand to the start position. Participants were required to maintain this position for 500 ms before the target appeared.

Targets (10-mm diameter) were displayed at one of several locations along a blue ring with a 70-mm radius. Upon target appearance, participants were instructed to perform a rapid, “shooting-style” movement to try to intersect the target with the cursor. Once the hand left the start position, continuous or delayed endpoint cursor feedback was provided depending on task conditions.

To enforce fast, ballistic movements, the maximum allowed movement time was limited to 300 ms. If the movement exceeded this limit, an auditory warning (“too slow”) was played. Such a speed limit for movement was deemed sufficient to prevent online correction of the movement (Poh and Taylor, 2019; Chen and Taylor, 2025). When the hand crossed the radial distance corresponding to the target location (70 mm from the start), a neutral “thunk” sound was delivered to indicate that the reach had terminated. Cursor feedback remained visible for 0.5 s following movement completion. Afterward, participants were guided back to the start location using the dynamic centering ring, and the next trial was initiated.

### Experiment 1

Participants (n=86) were randomly assigned to one of two groups designed to bias learning toward distinct cognitive strategies: an algorithmic strategy group or a retrieval strategy group (Figure 1B, C). Regardless of group assignment, the experiment consisted of baseline, training, and washout phases, totaling 484 trials (Figure 1E).

The baseline phase comprised three components intended to familiarize participants with the task, assess inherent movement biases, and ensure stable calibration before training began. First, participants completed a 16-trial familiarization block, during which the experimenter provided detailed instructions on performing the rapid center-out shooting movements and the goal of the task. Next, participants completed a 48-trial no-feedback block in which the cursor disappeared immediately after movement onset to obtain an unbiased estimate of each participant’s movement biases that may arise without feedback (Wang et al., 2024). Participants were instructed to aim directly toward each target and maintain fast, ballistic movements despite the absence of feedback. Finally, participants completed a 16-trial feedback block, during which veridical, continuous cursor feedback was restored. This brief block ensured that participants recalibrated the mapping between hand movement and cursor motion before entering the training phase. Across all baseline trials, cursor feedback – when provided – accurately reflected true hand position.

Following baseline, participants assigned to the retrieval group received five training trials at the Critical target with a 45° visuomotor rotation. All participants were told to counteract this rotation by re-aiming in the opposite direction by 45°. This brief exposure phase was designed to enable participants in the retrieval group to begin caching an explicit aiming solution for the Critical target, thereby reducing the discovery process (Tsay et al., 2024) and minimizing reliance on visuomotor mental rotation later in the experiment (McDougle and Taylor, 2019). Importantly, the delay endpoint feedback was provided during this period to minimize the implicit recalibration under a non-retrieval scenario. In contrast, trials were omitted from the algorithmic group, ensuring that algorithmic participants would rely primarily on an algorithmic (mental rotation–based) strategy during adaptation.

For each participant, a target was pseudo-randomly selected from a pool consisting of target locations from 0° to 330° in 30° increments. We designated this training target as the *Critical target* for probing implicit recalibration. The Critical target locations were yoked between the algorithmic and retrieval participants to ensure that target locations were matched across groups (Figure 1B, C).

The training phase consisted of reaches to 10 target locations, including the Critical target, spanning ±60° to ±180° around the Critical target in 30° increments (Figure 1B, C). Importantly, no training targets were placed within ±60° of the Critical target, thereby creating a gap around the Critical target for probing generalization while minimizing direct contamination from error-based learning (Figure 1B, C). The remaining nine targets were designated as Non-Critical targets (Figure 1B, C). Although implicit recalibration at the Non-Critical targets was not of primary interest, they served to establish the task context necessary to dissociate algorithmic and retrieval strategies and to equalize the statistics of reaches across the workspace.

In the algorithmic group, continuous cursor feedback with a 45° rotation was provided at all target locations (Figure 1B). Experiencing the rotation across multiple targets has been shown to pressure participants into relying on a visuomotor mental rotation strategy (McDougle and Taylor, 2019). In contrast, for the retrieval group, cursor feedback was withheld for all Non-Critical target locations and was only displayed while reaching the Critical target (Figure 1C). Restricting rotated feedback to only the Critical target encourages participants to retrieve a cached explicit aiming solution, rather than compute a rotation on each trial. Removing feedback at all Non-Critical targets further minimized opportunities for visuomotor mental rotation and helped equate inter-trial timing across groups and control for the movement variability across the entire workspace. Additionally, to promote use of the algorithmic strategy, the critical and Non-Critical targets were all displayed in the same color, minimizing the formation of a stimulus-response association based on color context (Figure 1B). By contrast, in the retrieval condition, the Critical target was presented in a distinct color from the Non-Critical targets, which was intended to strengthen the stimulus-response association specific to the Critical target. In brief, our study aimed to examine whether distinct strategies influence implicit recalibration at the Critical target. To promote the use of different strategies for adapting to the 45° rotation at the Critical target, we assigned distinct task instructions, feedback, and target colors to the Non-Critical targets in each group. Importantly, this design not only biased strategy use for each group, but also equated training at the Critical target and balanced movement variability across the workspace.

During the training phase, participants intermittently completed short sets of Exclusion trials, in which cursor feedback was withheld, and participants were explicitly instructed not to use any aiming strategy (Taylor et al., 2014; Werner et al., 2015; Maresch et al., 2021; t Hart et al., 2024). These trials were designed to probe both the magnitude of implicit recalibration and its generalization function around the Critical target. Specifically, seven probe targets spanning -45° to 45° in 15° increments, centered on the Critical target, were presented once per Exclusion block (Figure 1D). Thus, each Exclusion block contained 7 trials, ensuring that each probe target received one reach. The order of probe target locations was counterbalanced both within blocks and across participants; thus, we implemented a total of 7 Exclusion blocks during the training phase (Figure 1E). Immediately before each Exclusion block, participants made a single reach to the Critical target with rotated cursor feedback. This ensured that implicit recalibration was maximally engaged at the Critical target before implicit generalization was assessed. After completing the seven Exclusion trials, rotated cursor feedback was restored, and participants resumed practicing the full set of training targets in the workspace. To summarize the structure of this repeating “mini-block” within the training phase: Participants performed 40 reaches (except during the first block with 60 reaches) to the standard training targets –including the Critical target – then made one final reach to the Critical target with rotated cursor feedback before experiencing seven Exclusion trials to probe the magnitude and generalization function of implicit recalibration. This mini-block sequence was repeated periodically throughout the training phase for 7 times (Figure 1E). In each mini-block, 40% of trials were directed to the Critical target, whereas the remaining 60% were allocated uniformly across the Non-Critical targets.

A total of 356 trials were completed during the training phase, encompassing all training reaches and the seven Exclusion blocks (Figure 1C). The sequence of target presentations was pseudo-randomized by carefully controlling the intertrial intervals (ITIs), ensuring that the spacing between successive critical-target trials was consistent across participants. This approach ensured sufficient practice of the Critical target while preventing the formation of stimulus–response associations in the algorithmic group and avoiding excessive decay of implicit recalibration memory (Hadjiosif et al., 2023). Following the training phase, participants completed a washout phase that consisted of 48 trials. Within these trials, we ensured that each target location received three washout trials, given that 16 targets (i.e., 1 Critical target, 9 Non-Critical targets, and 7 probe targets (including the Critical target) for Exclusion trials) were used to construct the training phase. The task was paused at the transition between each phase of the experiment, during which instructional text was displayed on the monitor to inform participants about the upcoming trial structure (e.g., *“During the following trials, the cursor will be removed and will not be rotated. Please stop using any strategy and reach directly to the target.”*). Additional reminder text was also displayed whenever visual feedback was withheld (e.g., during Non-Critical targets and Exclusion trials) to further emphasize that participants should refrain from using any aiming strategy.

### Experiment 2

Experiment 2 (n = 40) followed the same general structure as Experiment 1, consisting of baseline, training, and washout phases; however, several key modifications were introduced to isolate the influence of strategies on implicit recalibration at the Critical target while minimizing the effect of plan-based generalization from training at the Non-Critical target locations. All participants were equally and randomly assigned to the algorithmic and retrieval groups.

We suspected there was a greater possibility that a broader breadth of the implicit generalization curve observed in the algorithmic group from Experiment 1 was attributed to an error “spillover” effect from reaching adjacent Non-Critical targets, where the implicit recalibration also occurred. To reduce the influence of practicing Non-Critical targets adjacent to the Critical target, the set of training targets was shifted further away from the Critical target. Specifically, all Non-Critical targets were sampled from ±90° to ±270°, creating a 90° gap (a 60° gap was used in Experiment 1) on each side of the Critical target. This larger angular separation ensured that participants did not perform repeated reaches near the Critical target, thereby minimizing the possibility that training at nearby Non-Critical targets would affect the implicit generalization profile centered on the Critical target (Figure 3A, B).

In addition, endpoint feedback for all Non-Critical targets was presented with a 1.5-second delay, a manipulation that has been shown to suppress implicit recalibration at those target locations, even when participants experience systematic cursor errors (Brudner et al., 2016). Thus, while the Critical target continued to support implicit recalibration through standard online rotated feedback, all Non-Critical targets were effectively prevented from accruing implicit recalibration (Figure 3A, B). This design enabled a direct comparison with Experiment 1, allowing us to test whether implicit recalibration at the Non-Critical targets produced spillover that inflated the implicit generalization function at the Critical target. At the same time, it provided a cleaner test of whether distinct strategies differentially shape implicit recalibration.

Aside from these modifications, the overall experimental procedure – including the baseline structure, the implementation of strategy conditions (algorithmic and retrieval), the Exclusion blocks used to probe implicit generalization, and the washout phase – remained identical to Experiment 1.

### Experiment 3

Experiment 3 (n = 38) followed the same overall structure as Experiments 1 and 2, including baseline, adaptation, and washout phases. In Experiments 1 and 2, the broader implicit generalization observed in the algorithmic group was likely driven by greater reach variability, which produced an error spillover effect consistent with plan-based generalization. To increase the robustness of control for this confounding factor (reach variability) and isolate the contributions of algorithmic versus retrieval strategies, Experiment 3 implemented a modified task design based on the error-clamp paradigm (Morehead et al., 2017). In general, the error-clamp paradigm refers to an experimental method in which feedback is artificially constrained to follow a preset trajectory or fixed error, regardless of the participant’s actual movement (Figure 4A), thereby allowing researchers to isolate implicit error-based adaptation while minimizing the influence of online performance correction (Morehead et al., 2017). Implicit recalibration elicited through error-clamp feedback exhibits the same qualitative characteristics as those observed in conventional visuomotor adaptation (Morehead et al., 2017). We leveraged this paradigm to fully control for any potential differences in error feedback that may inadvertently become linked via plan-based generalization when using an algorithmic strategy, which may increase movement plan variability, versus a retrieval strategy.

A Critical target was randomly selected from 12 possible locations (0° to 330° in 30° increments), with the frequency of each Critical target location matched between the algorithmic and retrieval groups. During the training, participants executed reaches to 9 different locations, spanning 45° to 165° away from the Critical target in 15° increments (Figure 4A-C). In addition, the angular space between the Critical target and the 45° location was used to define the probe targets for the Exclusion trials (see Figure 4D) to measure implicit recalibration and generalization. We trained to reach the 45° location (the Critical location) most frequently, consistent with counteracting the 45° rotations at the Critical target used in Experiment 1 and 2. The locations other than the 45° are denoted as the Non-Critical locations.

For the algorithmic group, all locations from 45° to 165° relative to the Critical target were invisible. Instead, after participants moved the cursor to the start position, an instruction text (e.g., *“Move toward 75 degrees”*) appeared at the top of the screen, indicating the required movement direction relative to the Critical target (see Figure 4A-C). This required them to perform visuomotor mental rotation to re-aim their movement away from the Critical target to the assigned location. After the completion of the movement, no cursor feedback was provided for any of these trials; instead, participants received coarsely-grained performance feedback:

*“excellent”* for errors ≤ 5°,

*“good move”* for errors between 5° and 10°,

*“fair move”* for errors between 10° and 15°, and

*“poor move”* for errors > 15°.

In contrast, for the retrieval group, we wished to prevent participants from performing visuomotor mental rotation throughout training. Thus, all Non-Critical locations were displayed as visible targets, and cursor feedback was withheld (Figure 4B, C). Participants were instructed to reach directly to these visible Non-Critical locations without using any additional aiming strategy. However, participants were instructed to reach toward the 45° location relative to the Critical target (no visible target at 45^°^), and cursor feedback was clamped in the opposite direction to enable implicit recalibration (see Figure 4A). To encourage caching a strategy for moving toward this 45° location, participants experienced five pre-training trials to allow participants to cache the correct aiming strategy for the 45° location.

To briefly summarize the design of Experiment 3, for both groups during training, the cursor feedback (if available) was error-clamped at 15° away from the Critical target, in the opposite direction of the required aiming direction (Figure 4A). More importantly, this clamped feedback was only used for the critical 45° location to maintain consistency with Experiments 1 and 2, in which participants learned to counteract a 45° rotation at the Critical target. In contrast, for all Non-Critical locations in both groups, no cursor or error-clamp feedback was provided (Figure 4B, C). Consistent with prior experiments, Exclusion blocks were included to assess implicit recalibration and generalization, with seven probe targets positioned around the Critical target at 15° increments from –45° to 45°. Aside from these modifications, the overall experimental procedure – including the baseline structure, the implementation of strategy conditions (algorithmic and retrieval), the Exclusion blocks used to probe implicit generalization, and the washout phase – remained identical to Experiment 1 and 2.

### Implicit Measures and Performance

Aftereffects, the hallmark of implicit recalibration, were assessed using Exclusion trials embedded throughout the training phase (Figure 1C). Each Exclusion block began with a top-up reach to the Critical location (45° angular direction for Experiment 3) to prevent decay of the adaptation (Hadjiosif et al., 2023). Within each block, targets used for probing implicit recalibration were presented in a randomized order and counterbalanced across blocks and participants (Figure 1B). Participants were instructed not to use explicit aiming strategies, but instead to reach directly toward each probe target. During these trials, both online cursor feedback and visual perturbations were withheld. The hand angle (the extent of hand drift away from the aimed target) at movement completion served as the measure of implicit recalibration. This same measure was also used to estimate the degree of adaptation on every trial throughout the task.

### Analyses Performance

Two primary dependent variables were analyzed: adapted hand angles and reaction time (RT). Hand angles were computed as the angular deviation between the final hand position at reach completion and the target location. These angles were used to quantify performance during training and to assess the magnitude and spatial extent of implicit generalization during Exclusion trials. We intentionally preserved the sign of the hand angle relative to the target location to avoid inflating the measure by taking the absolute hand angle. RT was defined as the interval between target onset and the moment the hand moved more than 5mm from the starting position. Prior work suggests that RT reflects the computational demands of planning, which differ between algorithmic and retrieval strategies (McDougle and Taylor, 2019; Velazquez-Vargas and Taylor, 2024).

To evaluate group- and time-related differences, we conducted a linear mixed model analysis with group (algorithmic vs retrieval) as the between-subjects factor and time (Early vs Late) as the within-subject factor for hand angle and RT for all experiments. More specifically, we conducted separate analyses for the Critical target and Non-Critical targets. Our primary aim was to investigate how strategy influenced implicit recalibration and generalization at the Critical target. Thus, the analysis focused on determining whether the comparable performance observed at the end of training was achieved through the use of distinct strategies. Although the Non-Critical targets were not the primary focus of this study, they were critical for reinforcing strategy use at the Critical target, as the desired performance across the Non-Critical targets was designed to support the engagement of different strategies. The adaptation performance for Non-Critical target locations was computed as the absolute error (absolute angular distance between the endpoint cursor and the target location). This at least ensures a fair numerical comparison of performance at the Non-Critical targets between the algorithmic and retrieval groups, given that the Non-Critical targets were associated with different task demands in each group. Early training was defined as the average value across the first training block, while late training was defined as the average value across the last training block. By conducting these analyses, we sought to verify that our paradigm successfully dissociated the algorithmic and retrieval strategies into two distinct groups, as evidenced by their differing RT profiles, while ensuring that both groups achieved comparable levels of adaptation. RT was used as an indicator of the cognitive strategy engaged during task performance. Based on prior research, we expected the algorithmic group to exhibit significantly longer RTs than the retrieval group, reflecting the greater computational demands of algorithmic planning (McDougle and Taylor, 2019). In addition, we compared hand angles and RT from the training with the baseline to validate that learning has occurred.

To assess whether the two groups exhibited comparable levels of adaptation prior to the Exclusion trials across all seven blocks, we conducted a two-way repeated-measure ANOVA on hand angle with group (algorithmic vs. retrieval) as the between-subjects factor and time (Blocks 1–7) as the within-subjects factor. Additionally, to verify that strategy use was withheld in the algorithmic group during the Exclusion blocks, we ran a separate analysis of RT measured during the Exclusion block using the same model structure. If strategy use was successfully withheld while reaching probe targets, the algorithmic and retrieval groups should show similar RT profiles across these blocks. Additional verification of successful strategy withholding was obtained by comparing reaction time (RT) during the Exclusion trials with that measured during training. The reduction in RT during the Exclusion trials, relative to training, suggests that participants disengaged the strategy.

To quantify and compare the magnitude of implicit recalibration across all Non-Critical target locations, we evaluated group differences relative to baseline performance and relative to the implicit recalibration measured at the Critical target. This analysis validated whether any residual implicit recalibration arising from Non-Critical locations was equivalent between groups. Controlling for this factor strengthens the interpretability of the present findings by confirming that group differences cannot be attributed to variations in implicit learning at Non-Critical target locations.

### Generalization Analyses

Generalization functions were estimated by fitting the endpoint hand angles from the seven Exclusion targets with Gaussian tuning curves (Tanaka et al., 2009; Brayanov et al., 2012; Poh and Taylor, 2019):

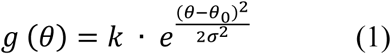

Where θ_0_ denotes the target location that produced the greatest shift in reach direction, k represents the amplitude of generalization, and σ characterizes the width of the tuning function. We used a bootstrapping procedure (1000 iterations), drawing from the population with exclusion and replacement, to estimate the generalization function (Poh and Taylor, 2019). This approach leverages the full distribution of samples (1000 total fits) and assesses parameter significance using 95% confidence intervals derived from the bootstrap distribution (α= .05; (Fisher, 1995)). Curve fitting was performed using MATLAB’s fmincon optimizer. Model fitting was conducted at the group level, based on each participant’s average hand angle at each probe target location across the 7 Exclusion blocks. Because implicit recalibration exhibits time-dependent decay, individual-level model fitting was not feasible unless the sequence of probe-target locations was fully counterbalanced across Exclusion blocks. Implementing such a design would have required 49 Exclusion blocks. Note, the hand angles from Exclusion blocks were normalized to the no-feedback baseline hand angles at the corresponding probe target locations before fitting the Gaussian model.

To control for potential plan-based generalization effects on the breadth of implicit generalization, we performed a separate analysis of movement variability at the Critical target location (Figure S6). Specifically, we quantified variability by calculating the standard deviation of the reaching directions for each bootstrap sample across 1000 resamplings. Greater variability in reaching direction suggests that participants generated more variable re-aiming plans. This dispersion in planned movement directions may broaden the resulting implicit generalization function (Day et al., 2016; McDougle et al., 2017; Chen and Taylor, 2025).

To demonstrate the relationship between reach variability and the breadth of the generalization function, we plotted the group differences in both the generalization function and the variability of endpoint hand positions (Figure 6A-B). The difference in the generalization function was defined, after model fitting, as algorithmic minus retrieval. Thus, positive values indicated greater implicit recalibration in the algorithmic group, whereas negative values indicated greater implicit recalibration in the retrieval group. Similarly, the difference in endpoint hand position variability was defined as algorithmic minus retrieval, such that positive values reflected greater reach variability in the algorithmic group and negative values reflected greater reach variability in the retrieval group. In Experiment 1, endpoint hand positions included reaches to both the Critical target and the adjacent Non-Critical targets, because implicit recalibration was present at those locations. In Experiments 2 and 3, endpoint hand positions included only reaches to the Critical target/location, because delayed endpoint feedback at the Non-Critical targets/locations reduced implicit recalibration and controlled for error spillover.

## Acknowledgements

We thank the members of the Intelligent Performance and Adaptation Laboratory (IPA Lab) for helpful feedback and insightful discussions. We especially thank Chandra Greenberg for providing comments on the manuscript. The research reported in this manuscript was supported by grant R01NS131552 (awarded to JAT) from the National Institute of Neurological Disorders and Stroke (NINDS) of the National Institutes of Health (NIH). The funder played no role in the study design, data collection and analysis, decision to publish, or preparation of the findings.

## Supplemental Information

**Figure S1.**
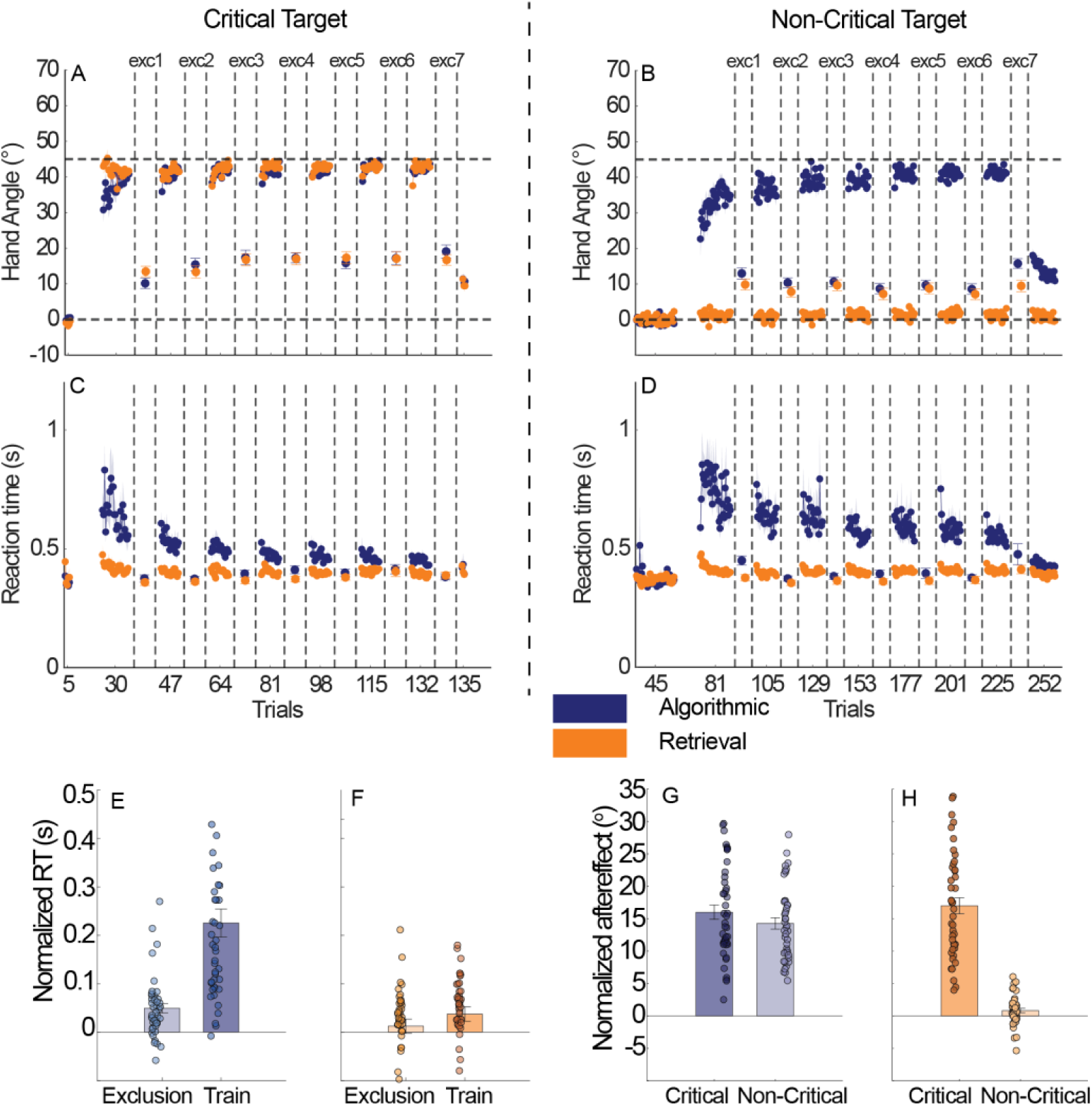
Full time course analysis for adaptation performance and reaction time, and control analysis in Experiment 1. A) signed hand angle and C) reaction time at the critical target. On the x-axis, the numbers indicate the cumulative trial count for reaches made to the critical target only. Dashed vertical lines mark the boundaries of the exclusion blocks (Blocks 1–7). Within each exclusion block, the error bars show the mean hand angle, reflecting implicit recalibration, and the mean reaction time measured at the critical target. B) Signed hand angle and D) reaction time at the non-critical targets. Here, the x-axis indicates the cumulative trial count for reaches made to the non-critical targets only. During each exclusion block, the error bars represent the average performance across the probe targets, excluding the critical target. E–F) Mean reaction time during Exclusion and training trials for the Algorithmic and Retrieval groups, also normalized to baseline performance. G–H) Mean hand angle during Exclusion trials at the critical and non-critical targets for the Algorithmic and Retrieval groups, shown as an index of implicit recalibration. These values were normalized to baseline performance.

**Figure S2.**
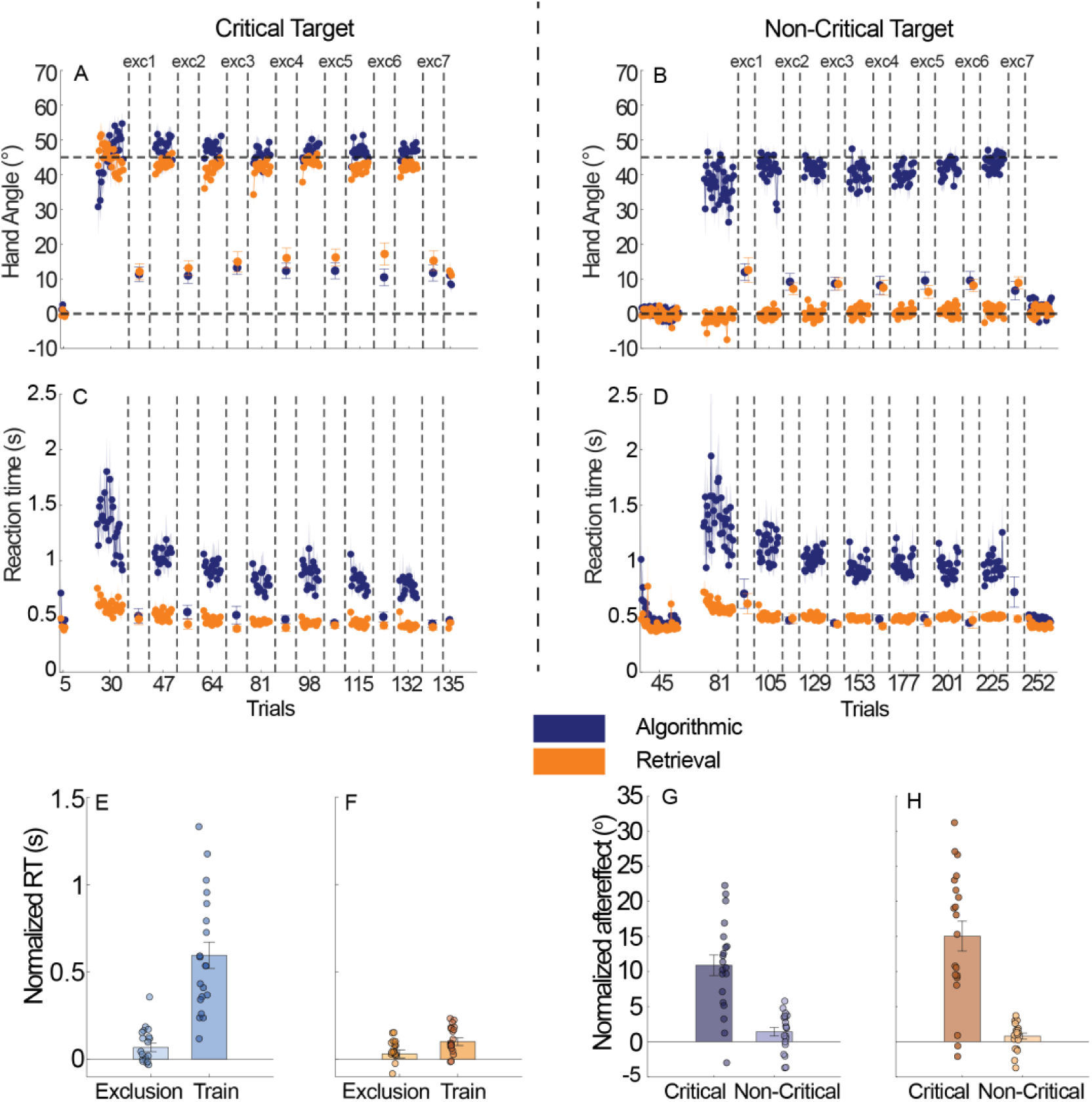
Full time course analysis for adaptation performance and reaction time, and control analysis in Experiment 2. A) signed hand angle and C) reaction time at the critical target. On the x-axis, the numbers indicate the cumulative trial count for reaches made to the critical target only. Dashed vertical lines mark the boundaries of the exclusion blocks (Blocks 1–7). Within each exclusion block, the error bars show the mean hand angle, reflecting implicit recalibration, and the mean reaction time measured at the critical target. B) Signed hand angle and D) reaction time at the non-critical targets. Here, the x-axis indicates the cumulative trial count for reaches made to the non-critical targets only. During each exclusion block, the error bars represent the average performance across the probe targets, excluding the critical target. E–F) Mean reaction time during Exclusion and training trials for the Algorithmic and Retrieval groups, also normalized to baseline performance. G–H) Mean hand angle during Exclusion trials at the critical and non-critical targets for the Algorithmic and Retrieval groups, shown as an index of implicit recalibration. These values were normalized to baseline performance.

**Figure S3.**
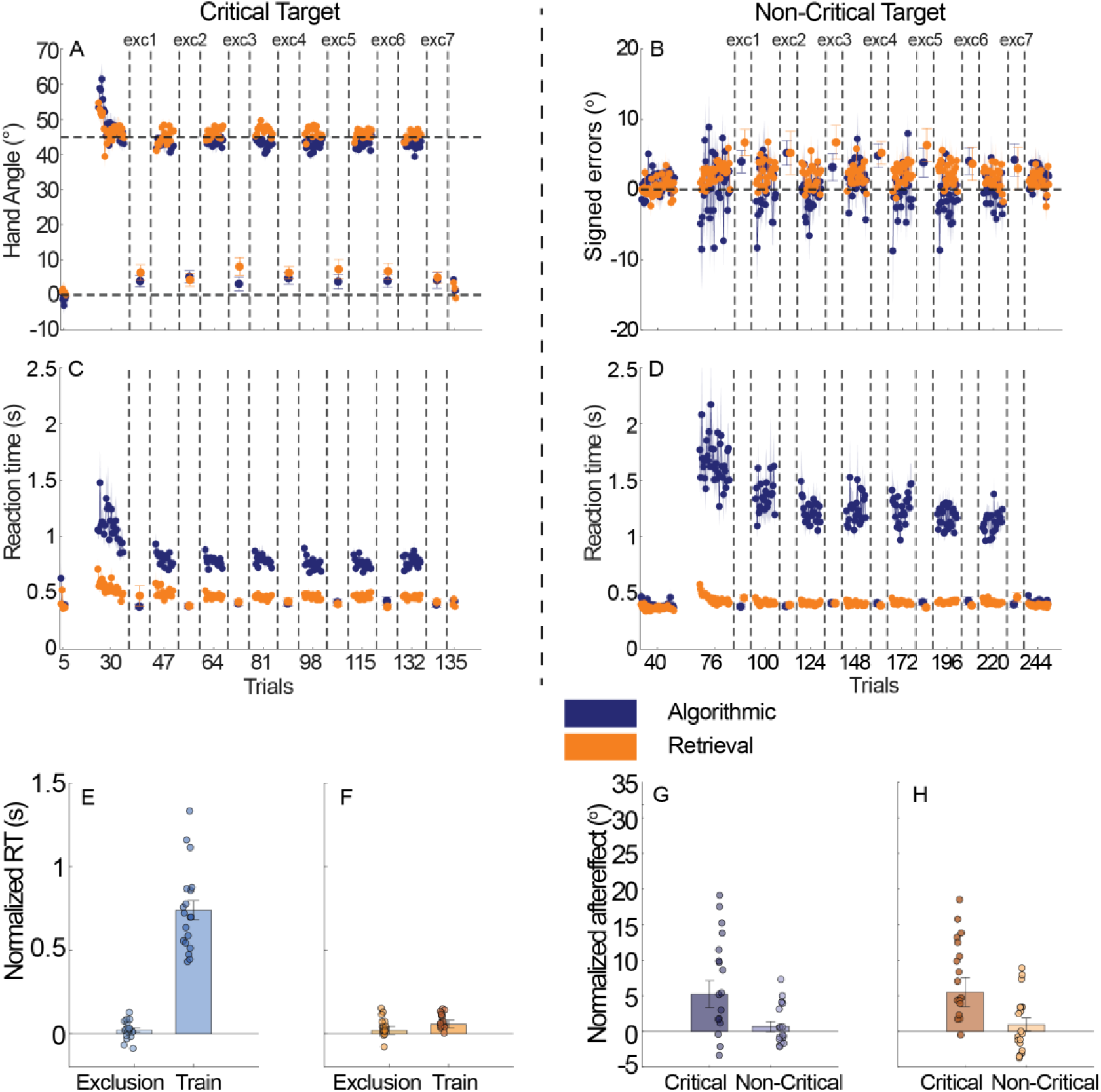
Full time course analysis for adaptation performance and reaction time, and control analysis in Experiment 3. A) signed hand angle and C) reaction time at the critical target. On the x-axis, the numbers indicate the cumulative trial count for reaches made to the critical target only. Dashed vertical lines mark the boundaries of the exclusion blocks (Blocks 1–7). Within each exclusion block, the error bars show the mean hand angle, reflecting implicit recalibration, and the mean reaction time measured at the critical target. B) Signed hand angle errors and D) reaction time at the non-critical targets. Here, the x-axis indicates the cumulative trial count for reaches made to the non-critical targets only. During each exclusion block, the error bars represent the average performance across the probe targets, excluding the critical target. The signed error is computed by subtracting the actual hand angle from the instructed reach angle. E–F) Mean reaction time during Exclusion and training trials for the Algorithmic and Retrieval groups, also normalized to baseline performance. G–H) Mean hand angle during Exclusion trials at the critical and non-critical targets for the Algorithmic and Retrieval groups, shown as an index of implicit recalibration. These values were normalized to baseline performance.

**Figure S4.**
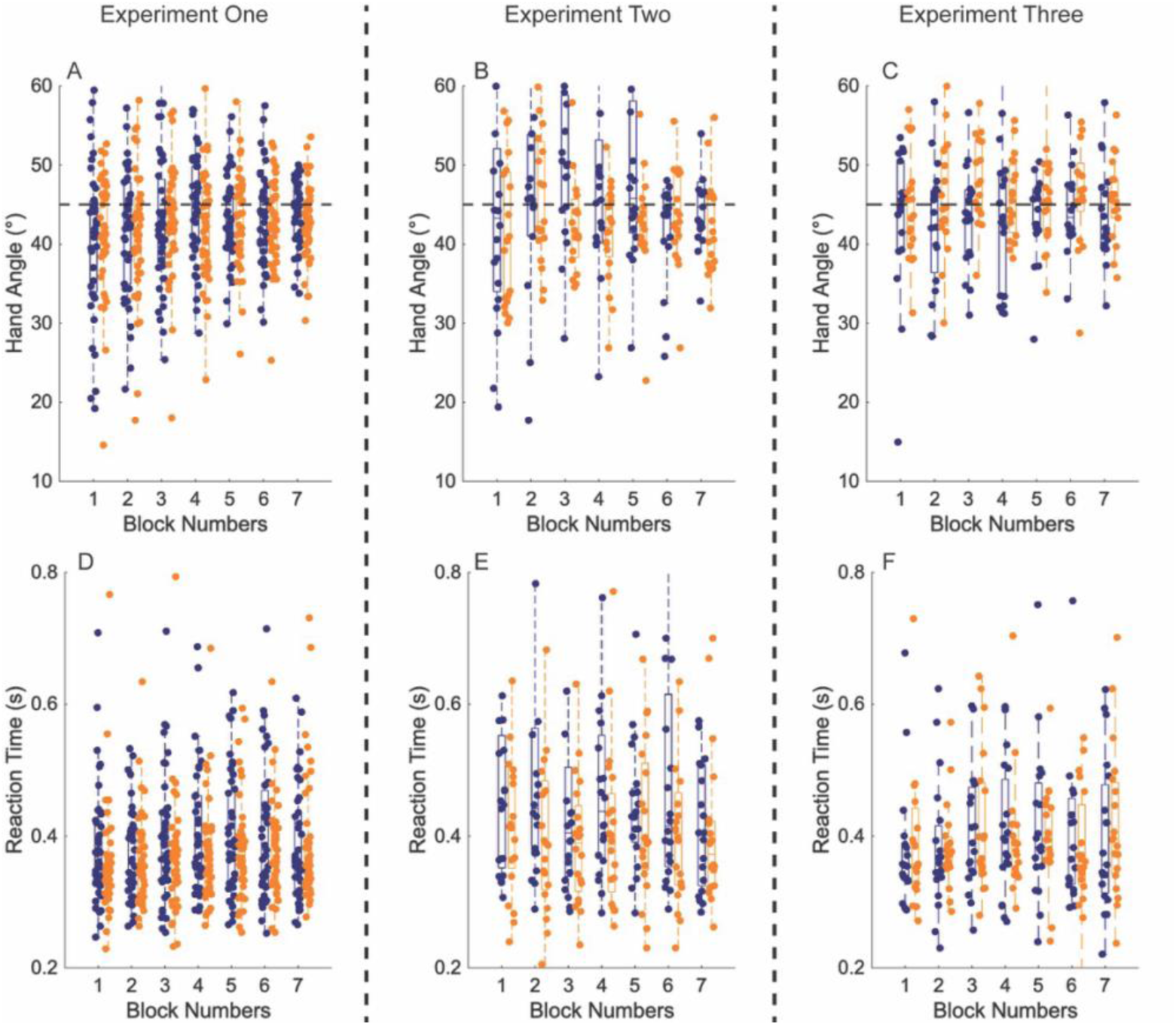
The extent of adaptation reflected by hand angles at a trial just prior to Exclusion blocks, and reaction time while performing during Exclusion trials. A–C) Hand reaching angles (°) for the single-trial performance at the critical target immediately preceding each Exclusion block across all seven blocks in Experiments 1, 2, and 3. These “top-up” trials were used to verify that both groups achieved comparable levels of adaptation before entering the Exclusion blocks, thereby preventing bias arising from potential time-dependent memory decay in implicit recalibration. The dash line represents the goal adaptation angle of 45°. As shown in figures, the algorithmic group (blue) and retrieval group (orange) reached comparable level of adaptation close to the goal 45° preceding each Exclusion block. D–F) Mean reaction times for all Exclusion trials within each block. During these trials, participants were instructed to withhold any strategy use, and thus the algorithmic group was expected to reach toward the probing targets as quickly as the retrieval group.

**Figure S5.**
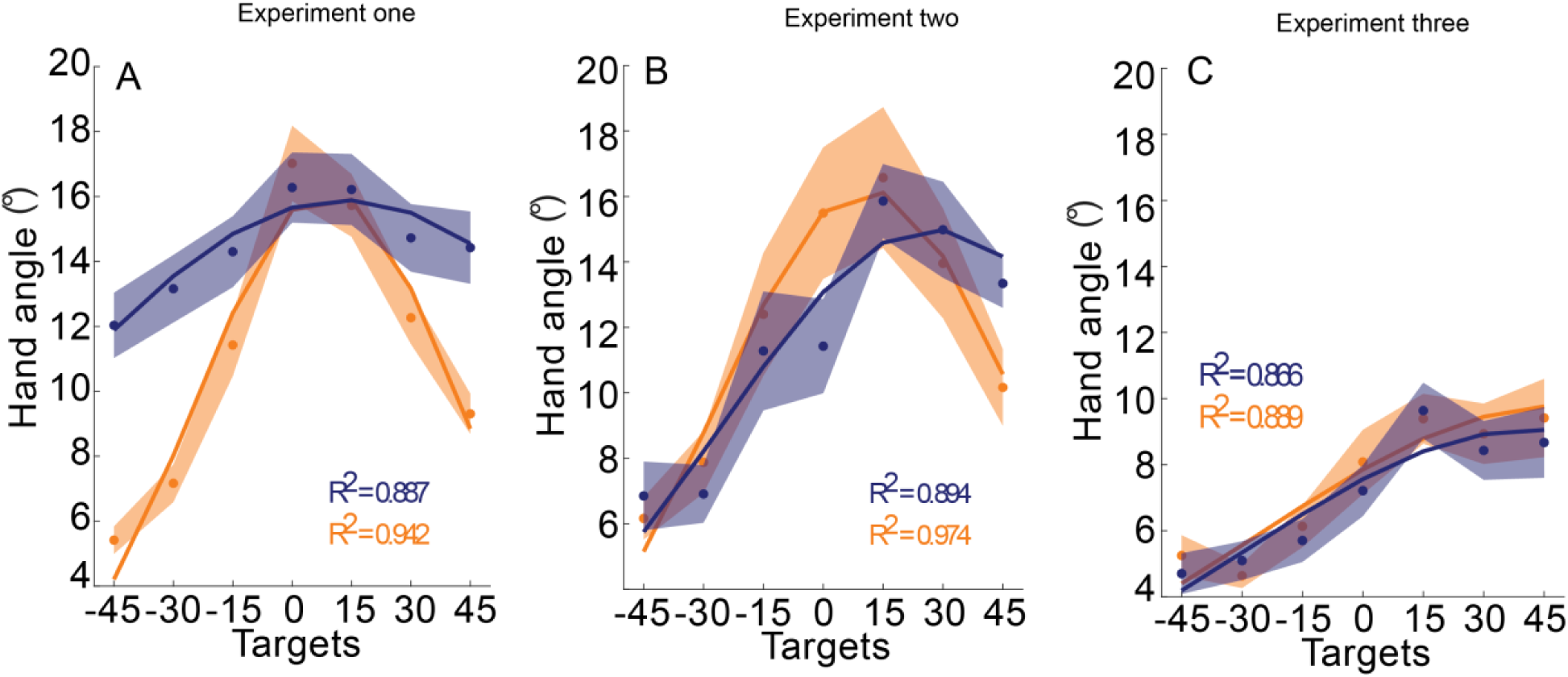
Gaussian fits for Experiment 1 – 3. A) The model fitting of the Gaussian function between hand angles and the 7 probe target locations (−45 ° to 45 °) in Experiment 1. The Algorithmic group is in blue, and the Retrieval group is in orange. B) The model fitting of the Gaussian function in Experiment 2. C) The model fitting of the Gaussian function in Experiment 3. Hand angles in all Experiments are normalized to the baseline.

**Figure S6.**
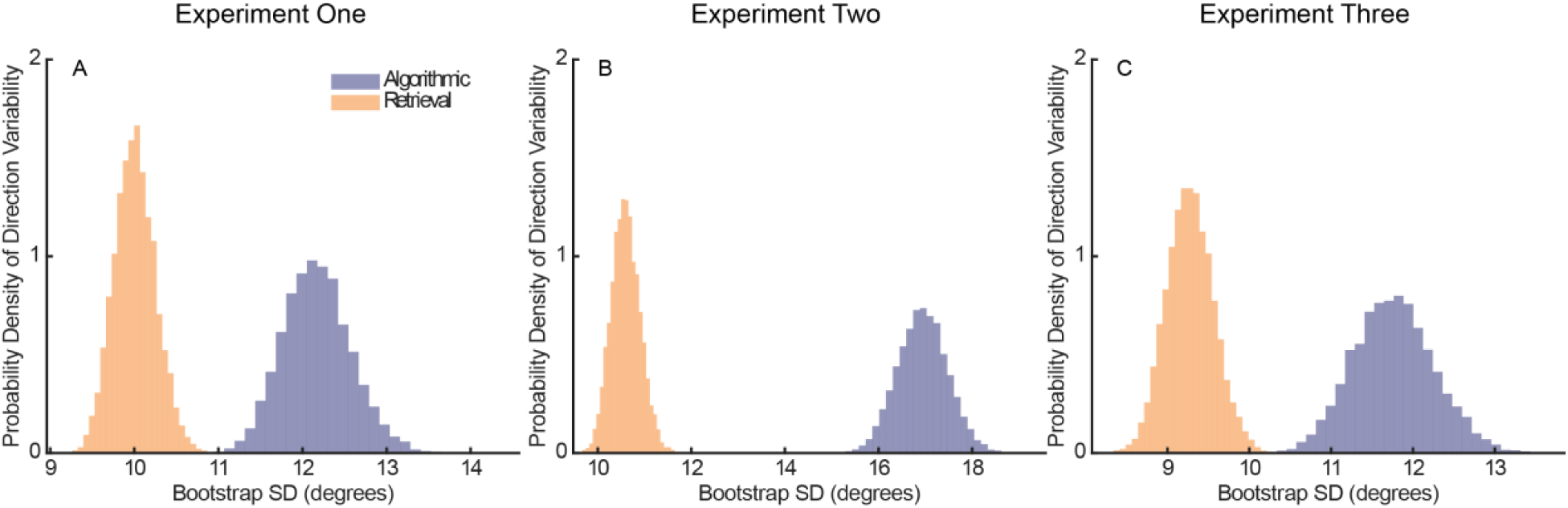
Distributions of reaching direction variability reflected by standard deviation. Implicit recalibration is thought to generalize around the planned movement direction, with each reach producing a small Gaussian-like kernel centered on that plan. Greater variability in reaching direction disperses these planned movement centers, so that learning accumulates across multiple nearby locations. The sum of these kernels broadens the overall generalization function across the workspace. To test this account, we examined movement variability at the critical target, reasoning that group differences in reach-direction variability could contribute to differences in the width of the implicit generalization function. Variability for each group was quantified as the standard deviation of endpoint hand angle at the critical target across 1,000 bootstrap resamples. In panels A–C, the x-axis shows the standard deviation of the endpoint hand angle relative to the critical target, reflecting the variability in participants’ reaching directions. (A) In Experiment 1, the Algorithmic group showed greater movement variability than the Retrieval group. Together with implicit recalibration arising from the non-critical targets, this increased variability likely contributed to a broader generalization function centered on the critical target. (B) In Experiment 2, the group difference in movement variability was even larger than in Experiment 1. Because implicit recalibration at the non-critical targets was minimal, variability around the critical target likely became the main factor underlying the broader generalization function. (C) In Experiment 3, implicit recalibration under error-clamp feedback was largely independent of aiming strategy. Although the Algorithmic group still showed somewhat greater movement variability, the implicit generalization functions were nearly identical across groups.

